# Demographic inference in a spatially-explicit ecological model from genomic data: a proof of concept for the Mojave Desert Tortoise

**DOI:** 10.1101/354530

**Authors:** Jaime Ashander, Peter Ralph, Evan McCartney-Melstad, H. Bradley Shaffer

## Abstract

In this paper, we study the general problem of extracting information from spatially explicit genomic data to inform inference of ecologically and geographically realistic population models. We describe methods and apply them to simulations motivated by the demography of the Mojave desert tortoise (*Gopherus agassizii*). The tortoise is an example of a long-lived, threatened species for which we have an excellent understanding of range, habitat preference, and certain aspects of demography, but inadequate information on other life history components that are important for conservation management. We use an individual-based model on a discretized geographic landscape with overlapping generations and age and sex-specific dispersal, fecundity, and mortality to develop and test a method that uses genomic data to infer demographic parameters. We do this by seeking parameters that best match a set of spatial statistics of genomes, which we introduce and discuss. We find that for inferring only overall population density and mean migration distance, a simple statistical learning method performs well using simulated training data, inferring parameters to within 10% accuracy. In the process, we introduce spatial analogues of common population genetics statistics, and discuss how and why they are expected to contain signal about the geography of population dynamics that are key for ecological modeling generally and conservation of endangered taxa.

## 1 Introduction

Mechanistic population models are a key tool used by basic and applied ecologists to understand the history and dynamics of natural populations. Population models inform fisheries management (Quinn and Deriso 1999), conservation of endangered species (Caswell 2001), and understanding of emerging infectious diseases (Diekmann and Heesterbeek 2000). Population models are well suited to address both fundamental questions (e.g., how population regulation occurs in spatially extensive, age-structured populations) and applied concerns (e.g., movement of disease vectors across political geographies).

Often, a population model is used to project future abundance under a variety of scenarios that affect one or more parameters (e.g., Prates et al. 2016; Benson et al. 2016). For example, a model that includes the effect of temperature on egg hatching rate could be used to project the impact of climate change on future population survival. To provide meaningful information for management, model parameters must be known with enough certainty that one can realistically distinguish the future effects of management or landscape modification scenarios.

Demographic quantities such as survival, growth, and fecundity can often be estimated by direct field observation (generally via mark-recapture studies) particularly for short-lived species where data can be collected over several generations. However, the degree to which current demographic parameters accurately reflect long term values is often unknown, particularly when those parameters may fluctuate substantially across time scales or geography. These demographic quantities determine abundance fluctuations and gene flow across the landscape, two processes with conceptually well-understood effects on patterns of genetic relatedness. Therefore, genomic data provide a promising source of additional information to bridge this gap. However, there has thus far been relatively little use of genomic data in fitting mechanistic ecological models, even though there must be a direct relationship between population dynamics and the geographical patterns of standing genetic variation observed in nature.

A major barrier to integrating genomic data in ecological models is a lack of analytical results that describe genetic patterns expected under geographically explicit population models. Genomic data are often used for descriptive models - most commonly, either clustering-based methods that seek to identify substructure in a population (e.g., Pritchard, Stephens, and Donnelly 2000; Bradburd, Coop, and Ralph 2017), “resistance” methods that depict genetic similarity using a landscape descriptor of gene flow (e.g., McRae 2006; Petkova, Novembre, and Stephens 2016; Shaffer et al. 2017), or least-cost path analysis to find most likely routes of gene flow (e.g., Wang, Savage, and Shaffer 2009). Although these approaches can provide information about migration rates among a set of discrete populations (Greenwald 2010), none of these methods provide estimates in units that are directly interpretable as describing population dynamics in a generative model of continuous space, such as mean distance traveled by dispersing individuals per year, or number of adults per square kilometer. It is extremely well-understood how demography determines genetic patterns in large, randomly mating populations (e.g., the Wright-Fisher model), but when realistic geography and its idiosyncratic effects are introduced, few analytical predictions are available (but see Ringbauer, Coop, and Barton 2017).

Simulations have proven useful in bridging this gap between ecological models and genomic data. A variety of simulation approaches can shed light on evolutionary and some ecological processes (Hoban, Bertorelle, and Gaggiotti 2012). Such studies generally use likelihood free (e.g., Approximate Bayesian Computation) approaches that require choosing summary statistics that describe the high-dimensional outputs (genomic or genome-like data). These simulation tools allow for inference of migration among complex, but discrete, spatially structured populations. For example Vallée, Luciani, and Cox (2016) used a general-purpose individual-based modelling (IBM) library to model the dynamics of 37 recombining markers across all human chromosomes during the Neolithic expansion among Southeast Asian islands. Alves et al. (2016) inferred short- and long-distance dispersal in Eurasian Neolithic expansions of humans using `SPLATCHE2` (which combines forward simulations of population sizes with coalescent simulations of genetic data, Ray et al. 2010). Similarly, Prates et al. (2016) inferred past demography of neotropical forest lizards, and S. E. Harris et al. (2016) inferred recent population structure (and correlating it with urbanization) in white-footed mouse (Peromyscus leucopus) in the Northeastern USA, both using `fastsimcoal2` (Excoffier and Foll 2011).

Here, we use an ecologically realistic, individual-based model to simulate whole genomes of a closed population across a heterogeneous landscape. By simulating across a range of parameters and comparing results from our model to genomes obtained from real populations, we can make inferences about the parameter values corresponding to these real populations. Actual population-scale landscape simulations of individuals with gigabase-sized genomes pushes the limits of current computational feasibility. However, our method is made computationally feasible with large population sizes by recent advances in simulation methods (Kelleher et al. 2018), that allow more rapid simulation and simultaneously record the genealogical relationships across generations for all individuals across the simulated landscape.

This study is motivated by the Mojave desert tortoise, *Gopherus agassizii* and the need to create realistic spatial models to guide its conservation. The species lives across much of the Mojave desert in the Southwestern USA, and is threatened due to a combination of habitat destruction, mortality due to human-subsidized predators (Kristan and Boarman 2003; Esque et al. 2010), disease (M. B. Brown et al. 1994), vehicle-associated mortality (W. Boarman and Sazaki 2006), and other factors (Berry 1986; USFWS 2011). A substantial body of ecological fieldwork now characterizes desert tortoise habitat suitability (K. E. Nussear et al. 2009; USFWS 2011), and several sizeable demographic studies have estimated sex- and age-specific mortality and fecundity (e.g., Doak, Kareiva, and Klepetka 1994; Karl 1998; Reed, Fefferman, and Averill-Murray 2009). However, certain aspects of tortoise life history - in particular, the effects of juvenile and long-distance dispersal - remain relatively unknown. Since dispersal-mediated gene flow should leave strong signals across the genome, we can reasonably hope that genomic data could inform a mechanistic understanding of tortoise movement across the landscape.

In this paper, we (1) Develop a landscape-scale individual-based model (IBM) simulation that maps ecological parameters to a population pedigree; (2) Introduce a general class of spatial population genetic statistics and motivate their use for inference problems such as those modeled here; (3) Develop a statistical method to estimate dispersal and population density by comparing patterns of relatedness on the landscape to simulations of expected relatedness; and (4) Use simulated data to show that our method can simultaneously estimate dispersal and density to within 10% percent of their true values, as long as the dispersal scale is not too large.

We then infer two parameters from data produced under the model used for inference, presenting a limited test of the method. This reflects a common scenario where a great deal is known about certain ecological and demographic processes and the goal is to add information from genomic data. We establish a geographically and ecologically realistic population model, develop methods to produce consistent discretizations of the model, explore a class of spatial genetic statistics, determine procedures to match these spatial statistics between datasets, and assess our statistical power across a range of model parameters.

The statistical problem we face here is an *inverse problem*, conceptually similar to estimating the migration rate between two randomly mating populations of size *N* by inverting the analytic relationship 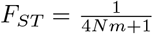(Wright 1951), where *F*_*ST*_ can be computed from genetic data, and *N* and *m* are the effective population size and migration fraction, respectively. However, the functional relationship between genetic statistics and parameters in models of continuous, heterogeneous geography is generally unknown (but see N. H. Barton, Depaulis, and Etheridge 2002; Ringbauer, Coop, and Barton 2017). Much like Approximate Bayesian Computation or sequential Monte Carlo (Marjoram 2013), we use simulation to bypass this issue. We simulate under a range of values of parameters for dispersal and density, calculate a large number of spatial population genetics statistics of the resulting data from each, and use general-purpose statistical learning to approximate the inverse map from statistics back to the (originally unknown) two parameters. In so doing, we demonstrate how spatially-explicit landscape genomic data can be used parameterize population biology models that should be useful across a wide range of applications and taxa.

## 2 Materials and methods

Although there are a great number of possible aspects of an ecological model that could be inferred, we focus here on a simple case. Suppose we have a set of of georeferenced whole genomes sampled from individuals across a population range, and that features of this spatial range have been recorded and interpreted to yield a measure of habitat suitability for the population of interest. However, both the rate of movement of individuals across geography (dispersal) and the population density remain unknown. We assume (i) that a local measure of carrying capacity is equal to *ρ* individuals per hectare, multiplied by local habitat suitability, and (ii) that the yearly movement of individuals is Gaussian with a standard deviation of *σ* meters. This leaves only two scalar parameters to be estimated: *ρ* and *σ*.

We model a landscape consisting of two large areas of high-quality habitat, which taper to low-quality habitat at their edges, connected by a narrow isthmus of low quality habitat (Figure 1A). Locations of individuals whose genomes have been sampled are marked by ‘+’ (in a real dataset, these would be fixed by the sampling location). From these individual samples, we seek to compute statistics that are informative of the population’s demographic parameters, *σ* and *ρ*.

**Figure 1:**
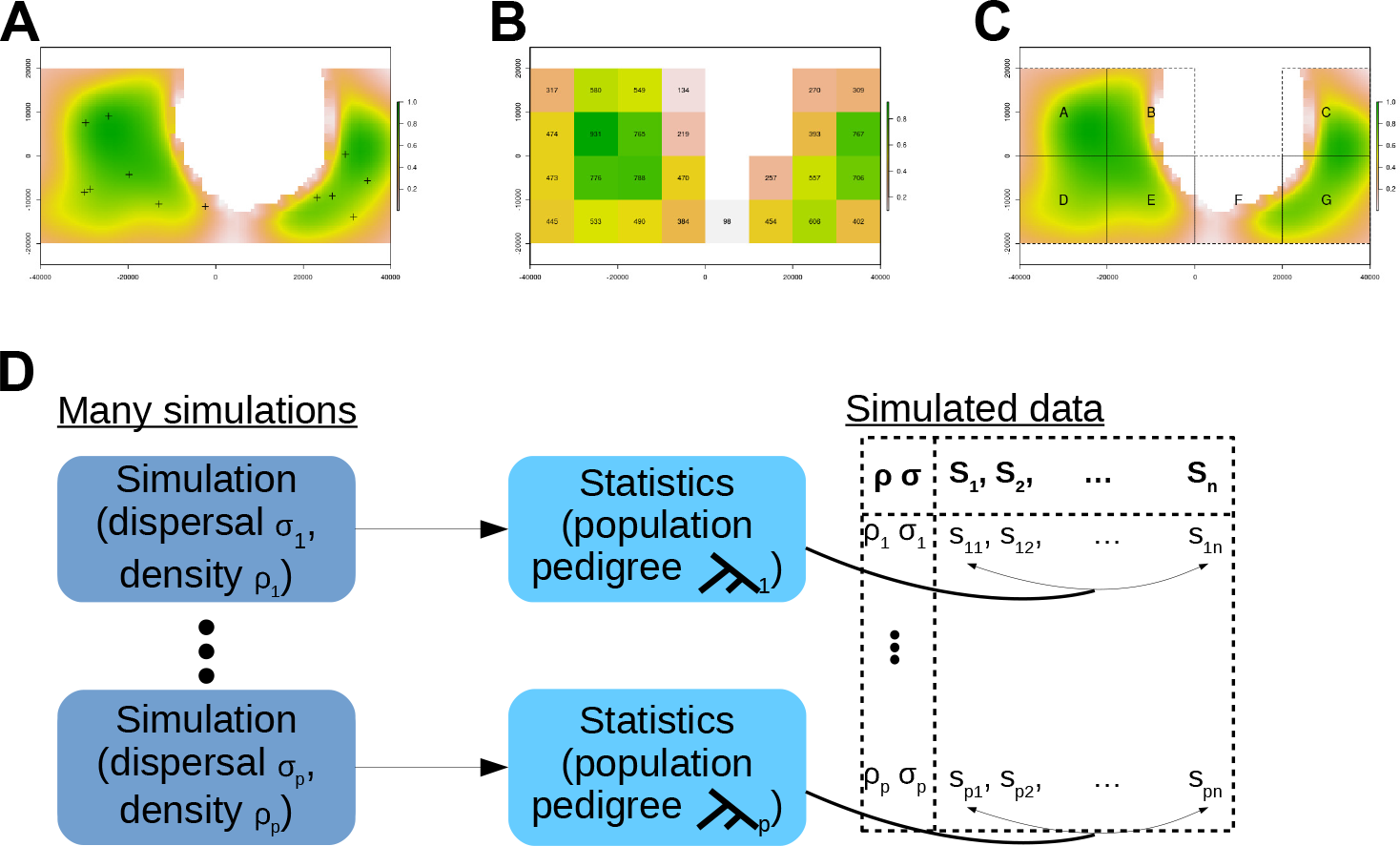
A) Spatial setting and example data, with axes labelled in meters and colors indicating habitat quality (1.0, or green, corresponds to highest-quality habitat, 0.0, or white, corresponds to impassable terrain) across the continuous geography. Individual samples are marked with ‘+’. For clarity only 12 samples are shown. B) The landscape discretized into patches used in simulations, defined by aggregating the fine scale map (panel A), and labelled with their carrying capacity of mature individuals (for *ρ* = 0.1). C) The landscape map partitioned into regions, labeled with letters; population genetic statistics are computed on groups of samples with group membership determined by the region in which samples occur. D) Simulating data for inferring dispersal *σ* and density *ρ*. We calculate *n* statistics for each of *p* independent simulations.

### 2.1 Geographic genetic statistics

To generate a wide class of potentially informative statistics, we use several population genetic statistics as *spatial statistics*, including Patterson’s F-statistics (Reich et al. 2009; Peter 2016). The *F*-statistics were originally used to compare variation among discrete, randomly-mating populations. In that context, the statistics convey information about admixture and shared branch lengths in the “population phylogeny” (Moorjani et al. 2013; Reich et al. 2009; Peter 2016) that describes how the populations are related to each other. Since we use these statistics in a nonstandard way, we now define them and motivate their use as informative spatial statistics.

For a set of genomes, denoted *A*, we write the *genetic diversity* of *A*, i.e., the mean density of nucleotide differences between a randomly chosen pair of genomes from *A*, as *π*(*A*) = 𝔼[|*a*_1_ − *a*_2_|], where *a*_1_ and *a*_2_ are alleles at a random site in the genome, coded as 0 or 1, from genomes randomly chosen without replacement from *A*. We also denote the *genetic divergence* between two groups, *A* and *B*, as the mean density of nucleotide differences between randomly chosen individuals from the two groups, which can be written as *π*(*A*, *B*) = 𝔼[|*a* − *b*|], where *a* and *b* now come from randomly chosen genomes in *A* and *B* respectively (if any individual is in both groups, sample without replacement so *π*(*A*) = *π*(*A*, *A*)). Patterson’s *F* statistics can be written in the same way, but using *four* genomes: *F*_4_(*A*, *B*; *C*, *D*) = 𝔼[(*a* − *b*)(*c* − *d*)], where *a*, *b*, *c*, and *d* are now alleles of randomly chosen genomes from the four groups *A*, *B*, *C*, and *D* respectively. Similarly, *F*_3_(*A*; *B*, *C*) = 𝔼[(*a*_1_ − *b*)(*a*_2_ − *c*)] and *F*_2_(*A*, *B*) = 𝔼[(*a*_1_ — *b*_1_)(*a*_2_ − *b*_2_)], where now *a*_1_ and *a*_2_ are randomly chosen without replacement from *A* (and likewise for *b*_1_ and *b*_2_). Note that we can write *F*_3_(*A*; *B*, *C*) = *F*_4_(*A*, *B*; *A*, *C*) if we interpret the latter as sampling without replacement, as we did for *π*(*A*, *B*); for this reason, subsequently we write all *F* statistics using this format (and dropping the subscript ‘4’).

We also introduce analogous statistics that depend on choices of *three genomes*. We will write these in terms of *y*(*A*; *B*, *C*) = 𝔼[*a*(1 − *b*)(1 − *c*) + (1 − *a*)*bc*], the probability that a sample from *A* differs from two other samples, one from *B* and one from *C*. The three-point statistics we use derive from *y*, but are modified to be zero in a randomly mating population: *Y*(*A*; *B*, *C*) = *y*(*a*; *b*, *c*) − (1/2)(*y*(*b*; *a*, *c*) + *y*(*c*; *a*, *b*)), and *Y*_2_(*A*; *B*) = *Y* (*A*; *B*, *B*) = *y*(*a*; *b*_1_, *b*_2_) − (1/2)(*y*(*b*_1_; *a*, *b*_2_) + *y*(*b*_2_; *a*,*b*_1_)).

An alternative way to think of these statistics is as estimates of weighted averages of branch length across all genealogical trees relating individuals selected from two or more groups, and scaled by mutation rate (Ralph 2015; Peter 2016). Averaging over marginal gene-trees under an infinite sites model of mutation, and omitting a scaling factor of the mutation rate, the corresponding “branch length” quantities (denoted with a superscript (b)) are:

- *π*^(b)^(*A*, *B*): the average sum of the lengths of the two branches going from *a* and *b* back to their most recent common ancestor (MRCA), averaged over trees and choices of *a* and *b*.
- *π*^(b)^(*A*): the same as *π*^(b)^(*A*, *B*) but with both genomes chosen from *A*.
- *Y*^(b)^(*A*; *B*, *C*): the difference between (the average length of any branches that separate *a* from *b* and *c*) and (one-half of the sum of the lengths of any branches that would separate either *b* or *c* from the other two), averaged over trees and choices of *a*, *b*, and *c*.
- *F*^(b)^ (*A*, *B*; *C*, *D*): the difference between (the average length of any branches that separate *a* and *c* from *b* and *d*) and (the average length of any branches that separate *a* and *d* from *b* and *c*), averaged over choices of *a*, *b*, *c*, *d*.

To avoid scaling factors and to make this correspondence exact, we measure branch lengths in *expected number of mutations*, i.e., scaling branches by the mutation rate per unit time. For instance, since *π*(*A*, *B*) is the average number of mutations per site that have occurred between *a* or *b* and their MRCA, this makes the expected value of *π*(*A*, *B*) equal to *π*^(b)^(*A*, *B*) under an infinite-sites model of neutral mutations. These relationships between statistics computed using genotypes and the summaries of branch lengths are shown in Figure 2.

The *F* and *Y* statistics are defined so that they have expected values of zero if samples are exchangeable (e.g., if they all come from a single randomly mating population), because in this case each topology is equally frequent and has the same distribution of branch lengths, so the contributions of each topology cancel. Figure 2 also shows formulas for the statistics in terms of divergence.

To help develop an intuition for how the statistics work in continuous geography, consider the situation where population density is constant, and movement in any direction is equally likely. Define groups of individuals by whether they fall in different geographic regions. Then, since regions of equal area have equal population size, distance between regions determines how closely connected they are by dispersal - i.e., how likely is a recent MRCA - analogous to the migration rate for discrete populations.

This makes it possible to consider the relative frequencies with which various marginal genealogies occur, and hence in which settings *Y* or *F* are expected to have positive, zero, or negative values. The frequency with which the first coalescence occurs provides a particularly good heuristic. For example, if group *A*’s region lies physically between the regions of groups *B* and *C*, then *Y*(*A*; *B*, *C*) will be negative due to a deficit of trees with topology (*A*, (*B*, *C*)), which has overall positive weight, as shown in Figure 2B. Conversely, if *B* or *C* lie between the other two, then *Y*(*A*; *B*, *C*) will be positive. Figure 3 shows several geographic configurations of regions and corresponding effects on *F* and *Y*. Across sites where none of the samples are closely related - e., there is no *recent* coalescence - all three rooted topologies in Figure 2B are roughly equally likely and have equal branch lengths on average (Wilkins 2004), and will therefore cancel out.

Population size also affects the probability of first coalescence. For example, *F*_2_(1, 2) = *F*(1, 2; 1, 2), which corresponds to Figure 2(C) with *A* = *C* = 1 and *B* = *D* = 2. If within-group coalescence is more likely than between-group (as is often the case), then unrooted topology (*AC*)(*BD*) = (11)(22) is the most common, implying that *F*_2_(1, 2) > 0. However, an increased population density in one region reduces the magnitude of *F*_2_, since it makes it more likely that two lineages in that region trace back to ancestors outside the region before they coalesce, decreasing this bias. Thus, increased population density is expected to decrease the value of the statistic. Figure 4 shows how varying region size (i.e., group population size) affects *F* and *Y*.

#### How many statistics?

If we have divided our samples into *k* groups we can compute a large number of statistics. The statistics we consider here are averages over choices of two, three, or four genomes chosen from varying numbers of groups. We call a statistic that depends on *k* groups a “*k*-point statistic” since we imagine each group standing in for a spatial location. “Groups” may be arbitrary: single genomes, single diploids, or larger collections. The total number of statistics of each type is:

- diversity, *π*(*A*) = *π*(*A*, *A*), a one-point statistic, *k*.
- divergence, *π*(*A*, *B*), a 2-point symmetric statistic, *k*(*k* − 1)/2.
- *F*_2_(*A*, *B*), a 2-point, symmetric statistic, *k*(*k* − 1)/2.
- *Y*_2_(*A*; *B*), a 2-point statistic, *k*(*k* − 1).
- *F*_3_(*A*; *B*, *C*) and *Y*_3_(*A*; *B*, *C*), both 3-point statistics symmetric in *B* and *C*, *k*(*k* − 1)(*k* − 2)/2.
- *F*_4_(*A*, *B*; *C*, *D*), a 4-point statistic with one symmetry and two anti-symmetries, *k*(*k* − 1)(*k* − 2)(*k* − 3)/8.

As the number of groups grows, it becomes impossible to compute all possible statistics in reasonable time. For example, with groups of size 20 each statistic takes 15 seconds to compute for a 10Mb region of a human genome (using Python tools from Kelleher, Etheridge, and McVean 2016), so for *k* = 30 it would take roughly 451 hours to compute all 108,345 statistics. For a fixed amount of computing power we must either keep the number of groups reasonably small and compute all statistics, or choose sets of statistics to compute, motivated by the biology or geography of the landscape.

#### Branch lengths or sequence?

The theory outlined above equates each statistic to a corresponding summary of branch lengths in marginal genealogies. Our simulations actually record all marginal genealogies that relate sampled individuals to one another at every point on the genome. As described in Kelleher et al. (2018), this is done for speed, but it has the benefit that we have access to the underlying marginal gene-trees. This means that we can directly compute expected values of the statistics on branch lengths. The alternative is to compute them using genome sequence, generated for simulations by placing mutations on the marginal genealogies. (Since neutral mutations do not, by definition, affect genealogies, placing mutations on the trees *post hoc* is equivalent to generating them as the simulation progresses.) For efficiency we take the former path, working directly with statistics calculated using the branch lengths of the underlying genealogies from simulations (e.g., using *F*^(*b*)^ rather than *F*). We expect performance with sequence-based statistics to be identical, because the deviation of the sequence-based statistic from the underlying tree-based statistic has mean zero with standard deviation inversely proportional to the square root of the sequence length (Ralph 2015) - in practice, they will be quite close for large data sets. A set of simple simulations confirms this predicted close agreement between the two methods (See Supplement B and Figure B.1). With empirical data, the statistics *must* be computed using sequence differences, but these are directly comparable to the branch length statistics after a rescaling.

### 2.2 A statistical method to infer population density and dispersal

Our overall goal is to estimate dispersal (*σ*) and population density (*ρ*) based on the statistics computed from a given genomic dataset with known geographical coordinates. Since the focus of this work is on the establishment of an ecologically realistic model and computation of informative spatial statistics, we do this in a relatively simple way. First, we simulate from an individual-based model across a grid of parameter values. Second, we compute statistics on each output. After this procedure (shown in Figure 1D) we obtain a table containing inputs (values of *ρ* and *σ*) and corresponding outputs, i.e., the statistics we calculate, denoted generically here as (*s*_1_ … *s*_*n*_). We seek to infer the relationship between inputs and outputs and do so using inverse interpolation.

**Figure 2:**
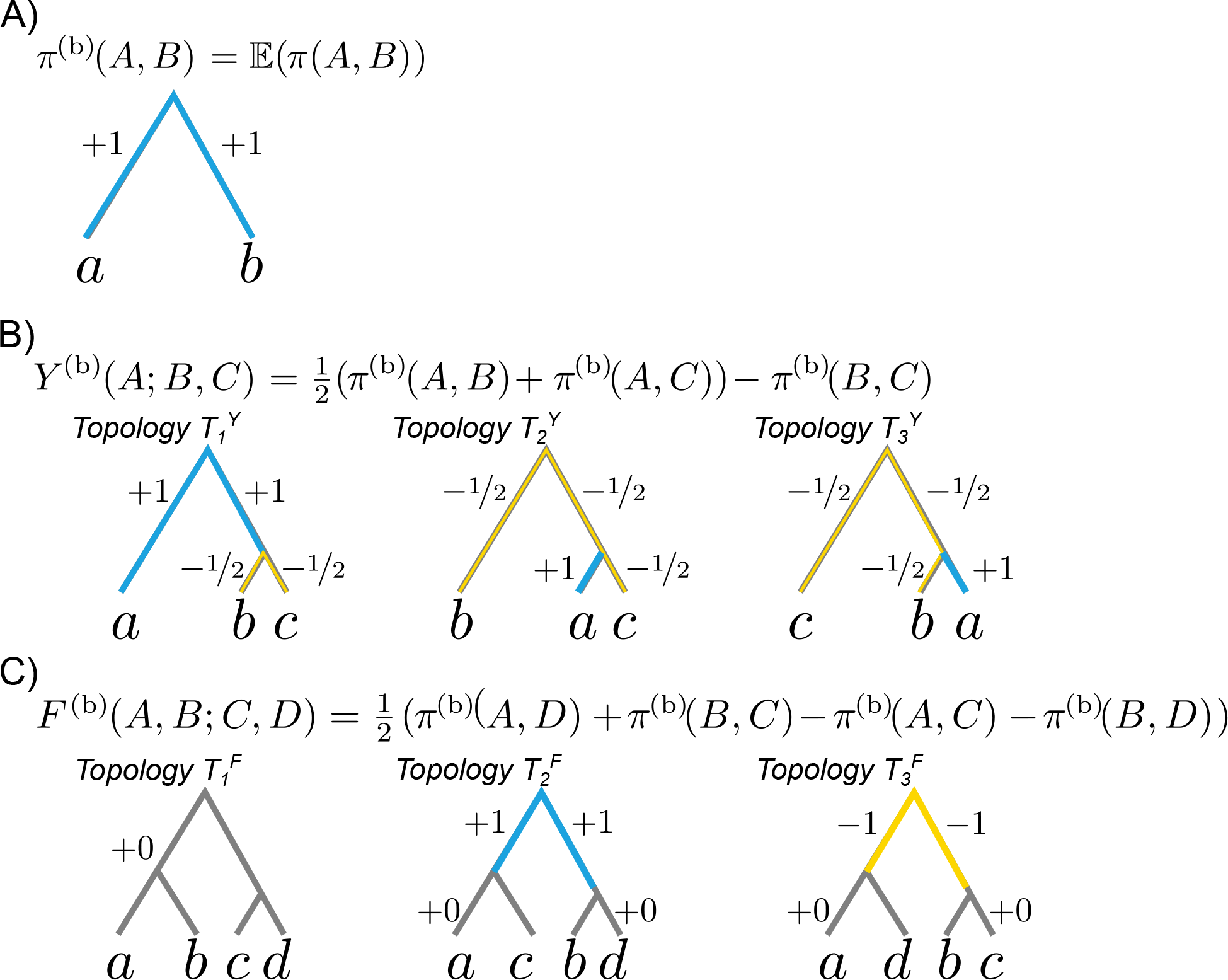
Genomic statistics between groups can be computed as a sum of weighted branch lengths across all gene trees; shown are the weights for the three types of statistic: (A) genetic diversity *π*, (B) the three-point statistic *Y*, the four-point statistic *F*. Branches are measured in number of mutations (for the usual statistics) or in units of expected mutations (for the ‘branch length’ versions). In the formulas, *π*(*A*, *B*) = *π*^(b)^(*A*, *B*) denotes the mean tree distance measured in terms of expected mutations from a random sample in *A* to a random sample in *B*, averaged over trees and choices of samples. Weights may depend on the tree topology and are marked on each branch; positive contributions are shown in blue and negative contributions are shown in yellow; gray is zero contribution. The equations in (B) and (C) show how the branch weights for *Y*^(b)^ and *F*^(b)^ in depend on those for *π*^(b)^ in (A). Equating *π*^(b)^(*A*, *B*) with the path between a sample from *A* and a sample from *B* yields the weights. For instance, in 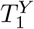 the weight of −1/2 on the branch above *b* (ii) is obtained by adding +1/2 (because it is on the path from *a* to *b*) to −1 (since it is on the path from *b* to *c*).

**Figure 3:**
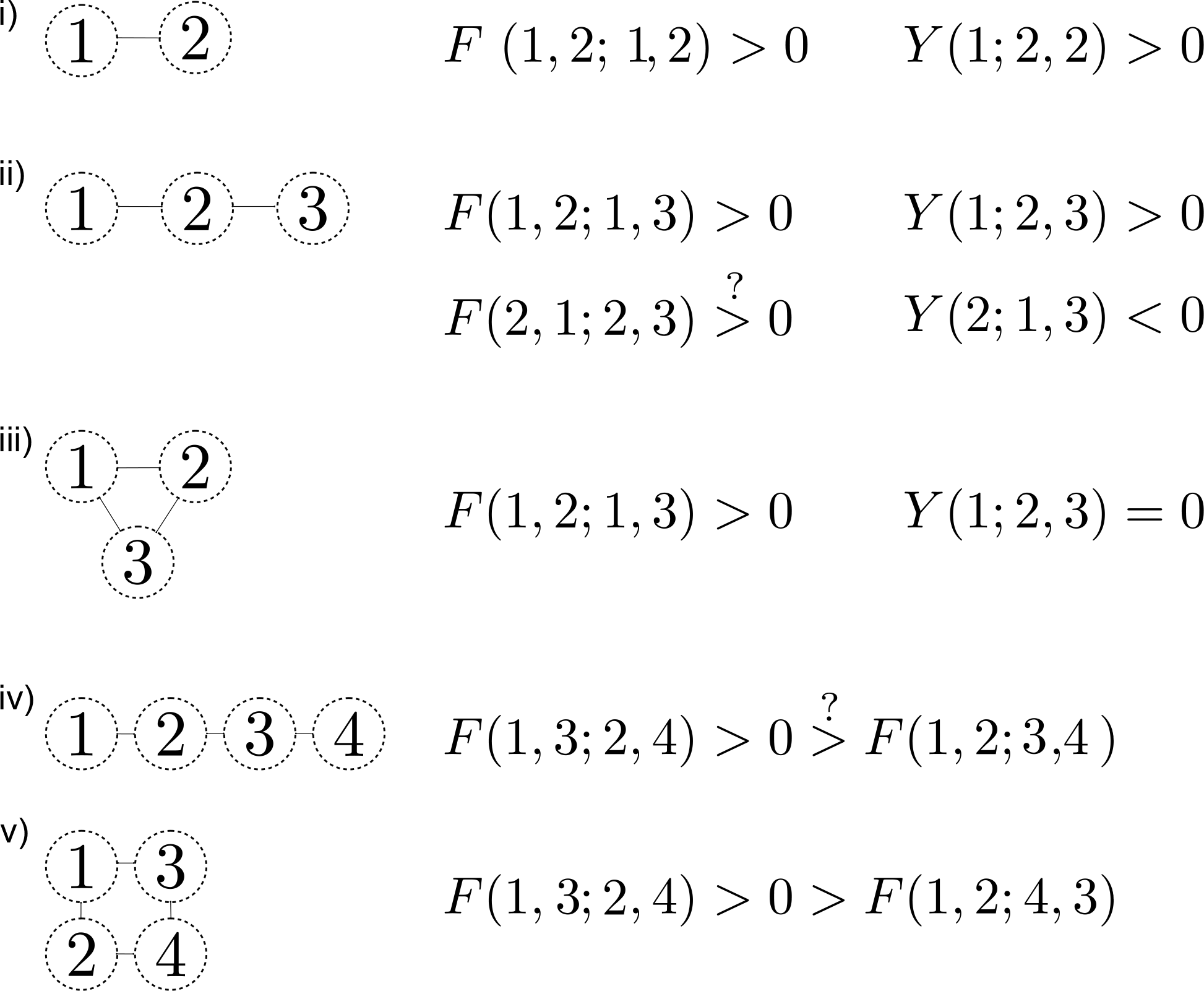
Effects of differing spatial configurations of regions in isotropic space on the signs of *Y* and *F* statistics. In all cases the sign is derived by reasoning about the probability of the first coalescence involving two regions and then connecting this to the probability that topologies occur in Figure 2. All population sizes (i.e., areas) are equal. Larger distances between regions in isotropic space result in decreased probability that the first coalescence involves these regions. Equivalent distances between equal-sized regions results in equal probability that the first coalescence involves these regions. If the actual sign is in doubt, the comparator is marked by ‘?’ Recall that *F*_3_(1; 2, 3) = *F*(1, 2; 1, 3) and that *F*_2_(1, 2) = *F*(1, 2; 1, 2). See Supplement A for justification.

**Figure 4:**
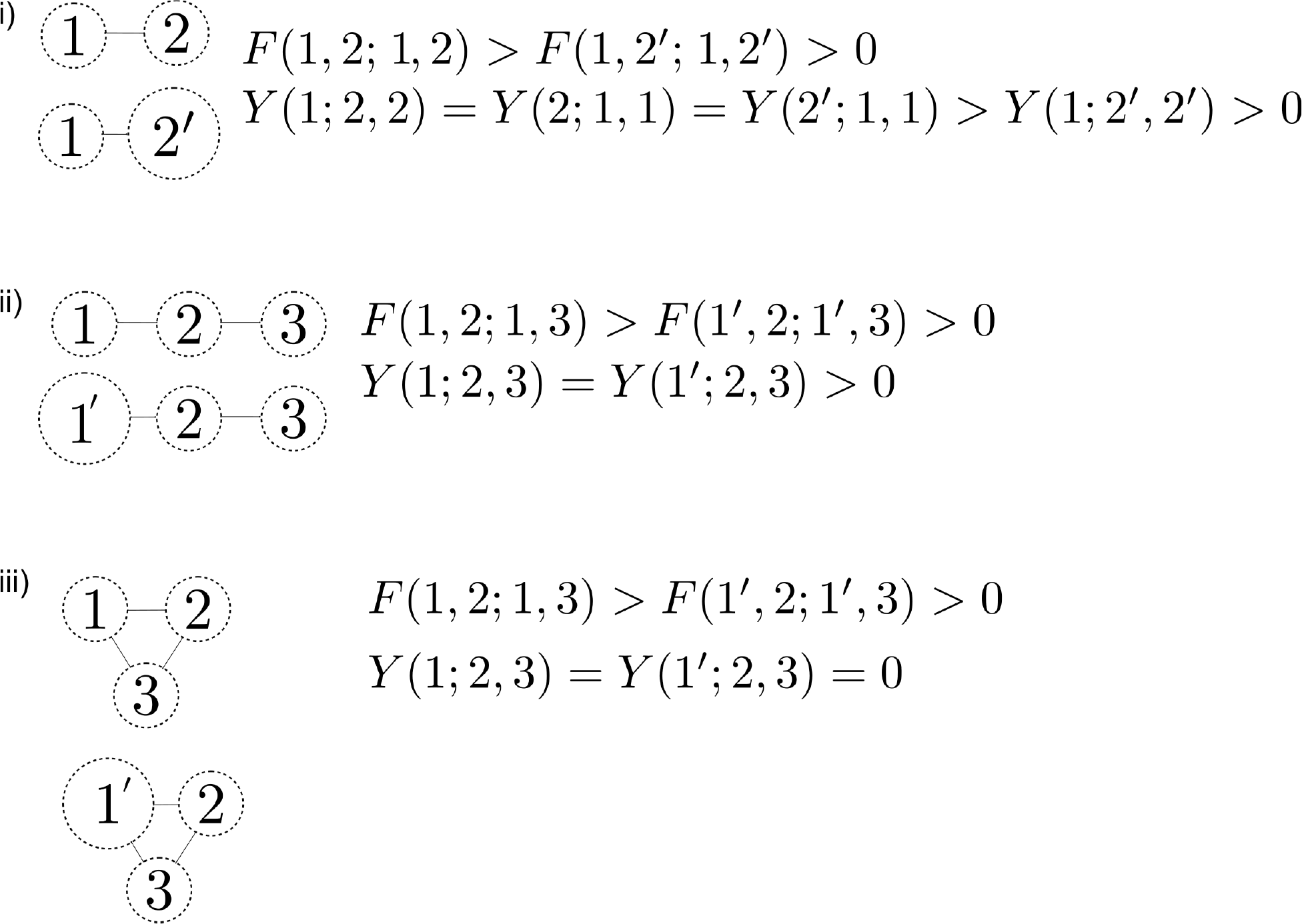
Effects of differing sizes and configurations of regions in isotropic space on *Y* and *F* statistics. In all cases the sign is derived by reasoning about the probability of the first coalescence involving two regions and then connecting this to the probability that topologies occur in Figure 2. All distances between centroids of adjacent populations are assumed equal. Regions denoted by larger circles have twice the population size (e.g., twice the population density) of those denoted by smaller circles. The probability of the first coalescence involving two members from a larger population is less than that of the first coalescence involving two members of a smaller population. Recall that *F*_3_(1; 2, 3) = *F*(1, 2; 1, 3) and that *F*_2_(1, 2) = *F*(1, 2; 1, 2). See Supplement A for justification.

#### 2.2.1 Inference via inverse interpolation

Consider using real genomic data to infer migration. In a classic Wright-Fisher model there is a clean parametric dependence of a genetic statistic, *F*_*ST*_, on migration rate *m* and population size *N*. For our more complex model there is an unknown analytic relationship between *σ*, *ρ*, and the statistics we introduce above. More generally, simulations give us noisy observations of an unknown function *f*(*θ*) that maps parameters (here, *θ* = (*σ*, *ρ*)) to the *n* statistics: for each simulation, run with parameters *θ*_*i*_, we can think of the resulting set of statistics as

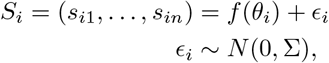

where Σ is an unknown covariance matrix defining how the noise *ϵ*_*i*_ is correlated across observations *i*. Given a new set of statistics *S̃*, we then seek to estimate the corresponding parameters, *θ̃*.

For our purposes, it suffices to take an average over the known parameter values *θ*_*i*_, each weighted according to the proximity of its associated statistics *S*_*i*_ to the observed *S̃*:

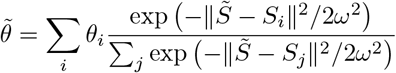

We choose the bandwidth, *ω*, by *k*-fold crossvalidation. To do this, we randomly divide the data *S* and *θ* into *k* blocks; denote by *S*^(*i*)^ the *i*-th block and *S*^−(*i*)^ the rest of the data. Then, for each bandwidth *ω* and each 1 ≤ *i* ≤ *k*, use the other data *S*^−(*i*)^ and *θ*^−(*i*)^ to predict parameters for every entry in the ith block *θ̃*^(*i*)^, and then compute the mean relative error in the *i*^th^ as:

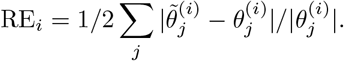

We choose the bandwidth *ω* that minimizes the median relative error across all *k* validation blocks.

#### 2.2.2 Individual-based simulations of a desert tortoise population

Our focal species, the Mojave desert tortoise (*Gopherus agassizii*) is widespread and exists as a collection of semi-discrete populations spread across the landscape. For species like this, one of the most important parameters for any spatial population model—dispersal—is also one of the most challenging to estimate empirically. In *G. agassizii* adults are generally dependent on burrows and therefore relatively stationary. But tortoises are also long-lived, so rare adult dispersal may provide significant, but rarely observed, genetic connectivity. At the same time, juveniles are secretive and have low survival, and movement is presumably much more common but also much more difficult to observe. Thus the long life, sparse distribution, and low juvenile survival rates all make it very challenging to accurately estimate the lifetime-averaged dispersal rate directly. Data from direct observations can provide some insights, particularly on short-term movement (e.g., radio telemetry, Nafus et al. 2017), but analyses of genetic samples from across the population’s range can potentially tell us much more about the recent history of movement in the population. The genomes of individuals encode information about their relatedness both across space and back through time. With genomic data from many individuals it is possible to accurately estimate statistics that describe these patterns of genetic relatedness and thus reflect movement and post-migration breeding of these elusive animals.

We developed a landscape-scale individual-based model (IBM) simulation of a closed population of interbreeding individuals, including relatedness across the genome. The model includes several demographic complexities, including sex (individuals are diploid, with sex fixed at birth), age structure (i.e., varying survival and fecundity by age class), density-dependence, movement in space, and maturity (which increases survival). Our IBM simulates a closed population, which could be an entire species.

We implement the model by discretizing continuous space into a grid of discrete patches, where the patches are contiguous and each represents a subpopulation (Figure 1B). In the simulation below, each of these includes approximately 50-500 individuals. Within each patch, we assume that individuals mate randomly. We also implement population regulation within the patches.

Movement between patches is parameterized by the standard deviation of the dispersal kernel, *σ*. We compute the *migration matrix M* whose (*i*,*j*)th entry is the probability that an individual in patch *i* moves to patch *j* in a given year. This is computed as the probability that an individual uniformly located in patch *i* moves a random, Gaussian-distributed distance with mean zero and standard deviation and ends up in patch *j*; if the corresponding geographic regions are denoted *A*_*i*_ and *A*_*j*_ then this can be computed as

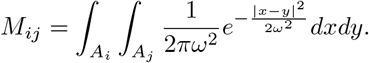

This computation is done by numerical integration using the **R** package `landsim` (Ralph 2017).

Our simulation performs one step in the **life cycle** for each *year*, as follows:

- Dispersal: Females each choose to disperse with probability 1/2; males are more vagile and disperse every year. Each dispersing individual in patch *i* independently chooses a new location, moving to *j* with probability *M*_*ij*_. Since *M*_*ii*_ > 0, this may result in no movement.
- Maturation: Newly born individuals are immature, and to mature they need to find available resources. The probability per immature individual of maturing is *K*/(*S* + *K*), where *S* is the local number of already mature individuals, and *K* is the local “carrying capacity”. Each subpopulation’s carrying capacity is equal to the product of *ρ* and the integral of habitat quality over the corresponding geographic patch.
- Birth: If there are available mates, every mature female produces offspring in a *single clutch*, mating with a randomly chosen male of reproductive age (at least 15 years old) from the same population as the mother, if any males are available. (If none are available, she produces no offspring.) The new offspring have age 0, and the number of these produced per clutch is Poisson with a mean that depends on age (see Supplement D), derived from (Reed, Fefferman, and Averill-Murray 2009).
- Growth: increment all ages by one year.
- Survival: kill individuals (including new ones) with probability depending on their age determined as in (Reed, Fefferman, and Averill-Murray 2009) and listed in Supplement D, Table D.2.

As our model structure is inspired by *G. agassizii*, parameters for survival and fecundity are both drawn directly from literature on tortoises. Potentially critical aspects of tortoise biology, however, are missing from the model including variation in dispersal rates by age.

Further details and parameters of the model are given in Supplement D.

##### Implementation with pedigree recording

We implemented our IBM in `simuPOP` (Peng and Kimmel 2005), a flexible individual-based simulation library with a python interface. Patches were implemented as subpopulations, with sex structure to allow different dispersal probabilities for males and females. Age-dependence of survival and fecundity were both implemented via `python` functions passed to `simuPOP`.

To record the population pedigree (actually the embellished pedigree recording all relationships between all genomic segments in the entire population, or ‘nedigree’) from our forward simulations into a ‘tree sequence’ data structure we used the efficient pedigree recording method described in Kelleher et al. (2018) and implemented in `python` for `simuPOP` in `ftprime` 0.0.6rc. In this method, haploid genomes correspond to nodes in the tree. Using data from every recombination, we record the tree structure on every genomic interval. Periodically during a forward simulation, this data structure is simplified, and its size reduced in memory, to the genomes of living individuals and their ancestors. At the end of the simulation, this tree sequence can be queried to describe underlying genealogies or genome sequence, and enables extremely fast computation of statistics.

### 2.3 Evaluating the method

We simulated single chromosomes of length 10^8^ base pairs, with a recombination rate of 10^−8^ per base pair per generation on the landscape of Figure 1A. We ran one simulation for 15,000 years at each of the 225 parameter combinations from 15 values of *ρ* evenly spaced in the range 0.05 to 0.2 individuals per hectare and 15 values of *σ* logarithmically spaced in the range 10 to 1000m per year. The range of density *ρ* corresponds to 500-2000 individuals in a patch with optimal habitat (value 1.0 on the landscape of Figure 1A). Although historic estimates of tortoise density are elusive (USFWS 2011), desert tortoises have been reported at densities on the order of 10 per km^2^ (0.1 per hectare) (Allison and McCoy 2014), and yearly home range sizes span 0.01-88.6 hectares (since 1980 in CA/NV from Table 11.1 of Berish and Medica 2014). These ranges are consistent with our ranges of *σ* and *ρ*. Simulations were initialized with the results of a coalescent simulation in a very small population, as for instance might result from a rapid expansion. This means that the populations are *not* at demographic equilibrium (at least for larger population sizes), which makes the inference problem both more difficult and more realistic.

#### Geographic regions for statistical computation

We merged adjacent patches into seven regions based on geography (Figure 1C). For each simulation, we sampled 500 individuals (see next paragraph for how these individuals are chosen) and grouped them according to the seven regions, so that individuals whose location fell in a region were all in the same group. We then computed all statistics between these seven groups of individuals By using only seven groups, we can compute all 406 possible statistics among the groups in a reasonable amount of time.

#### Choice of sampled individuals

Our next step was to compare statistics computed from each simulation to those obtained from data. However, these patterns of statistics can be sensitive to the geographic positions of the sampled individuals, even within the geographic regions defining each group. To remove this source of noise, we developed a scheme to choose, in each simulation, a set of individuals closely matching the spatial positions in our dataset. To achieve this, we first defined a reference simulation (with values *ρ* ≈ 0.104 and *σ* ≈ 100), and chose 500 individuals from the reference simulation. (For inference from non-simulated data the reference would be the empirical samples.) Our goal was then to choose individuals in other simulations that are geographically close to these. Suppose, however that we chose 50 reference individuals from a given patch; we are not guaranteed in a different simulation to have 50 individuals (total) in that same patch, a problem that becomes worse as the spatial discretization becomes finer. Therefore, for each reference individual we assigned weights to each patch corresponding to the distribution of that individual’s location after 800 migration steps (by applying the migration matrix 800 times to a vector that indicates the initial location). This yields 500 weightings of the patches to sample from, each corresponding to a reference sample. To choose individuals from other simulations, then, we choose a patch based on the weighting and then sample an individual uniformly from it. (If it is empty or all individuals have been sampled, we choose from the next patch.) This process is illustrated in Figure 5.

#### Evaluation by crossvalidation

We evaluated the model using *k*-fold crossvalidation. This is a similar procedure to how we chose 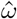 but now we use it to evaluate the model’s performance in terms of relative error. We randomly divided the simulated data into validation blocks. Then for every block we used the simulations not in that block to infer parameters for each simulation in that block. For each validation block we computed mean relative error. We did this using all possible statistics (*n* = 406): each of the six types computed across all seven groups.

In many other supervised learning contexts, having a large number of predictors (here, the statistics) relative to observations (here, the number of simulations) can cause overfitting and biased predictions (Hastie, Tibshirani, and Friedman 2009). Thus, for comparison, we did the same thing with a much smaller set (*n* = 7) of transformations computed on individual or groups of the statistics. These were designed based on the geographic and biological setting to give us more nearly independent information. We refer to these as “custom” or “biologically-motivated” statistics (see Supplement C for definitions of these).

**Figure 5:**
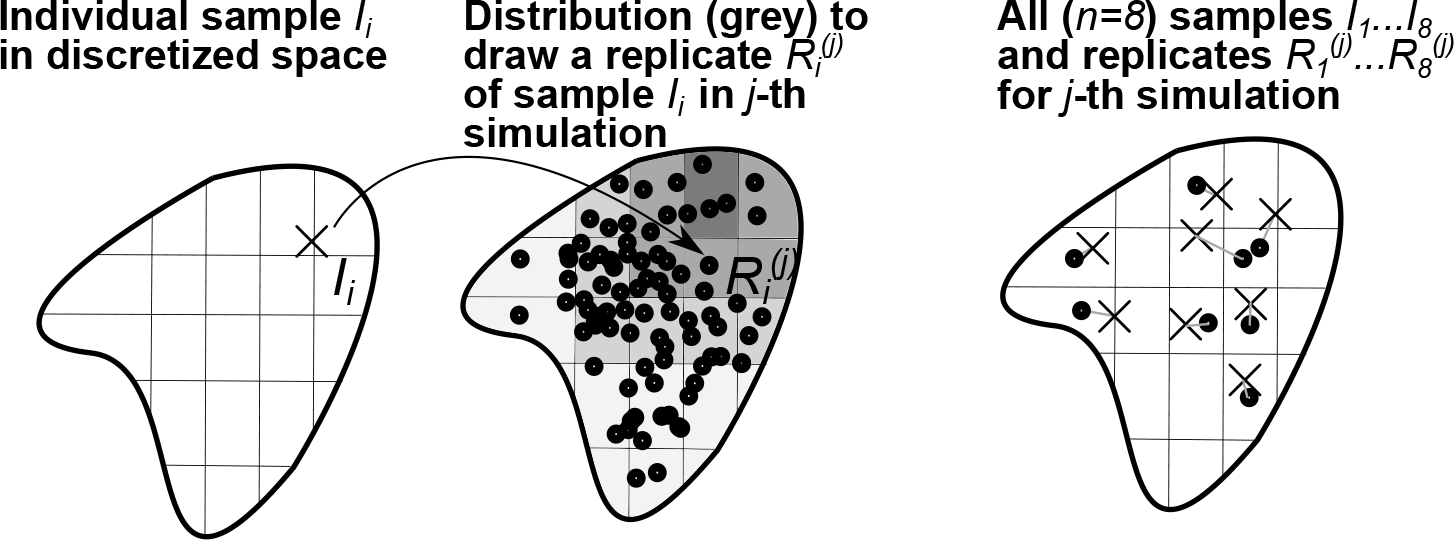
A method to choose simulated samples that match *n* reference samples: 1. identify patch locations of samples (*I*_*i*_) on the discretized landscape, 2. for each sample *I*_*i*_ construct a distribution to draw a replicate in the *j*-th simulation 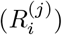 based on the location of *I*_*i*_ (solid arrow), 3. for each of *j* simulations draw the *n* replicates (connected by gray lines). Samples *I*_*i*_ could be empirical data or individuals in a reference simulation, in which case their location is simply their patch.

## 3 Results

### 3.1 Model behavior

The IBM simulations produced a population with age-structure and dynamics similar to that seen in the models of tortoise demography (e.g. Reed, Fefferman, and Averill-Murray 2009; Doak, Kareiva, and Klepetka 1994) from which the age-structure in the model was largely parameterized. For example, the shape of mean realized lifetime fitness (number of offspring) versus age for females (see Supplement D, Figure D.4) agrees qualitatively with that of reproductive value seen in other models.

### 3.2 The simulated data

Plots of all 406 statistics are difficult to interpret (see Supplement E). However, it is clear that many of the statistics carry significant signal regarding both density and dispersal. For example, separating relative divergence comparisons between regions into three categories — within regions, between neighbors, and between non-neighbors (Figure 6) — results in interpretable patterns. In self-comparisons, divergence dips well below the mean as dispersal declines, especially for less-dense populations. Meanwhile comparisons between neighbors display varying patterns that relate to geography. For example, the E-F divergence, which involves neighbors on the low-quality habitat isthmus (Figure 1A), behaved differently than the other statistics. Further intuition can be gained from examining other biologically-motivated combinations of statistics (see Appendix C, Figure C.2).

### 3.3 Performance

Using all possible statistics, the median relative error across all 5 cross-validation blocks was 0.104 (10.4%) with standard deviation of 0.016. In contrast, using only the few biologically motivated “custom” statistics the median relative error was 0.169 with standard deviation of 0.028. These results indicate the work needed to develop and specify statistics for a given geography was counterproductive: using only “custom” statistics had almost twice the prediction error as the method using all statistics. Figure 7 shows predicted versus actual values for the inferred parameters, density and dispersal scale.

**Figure 6:**
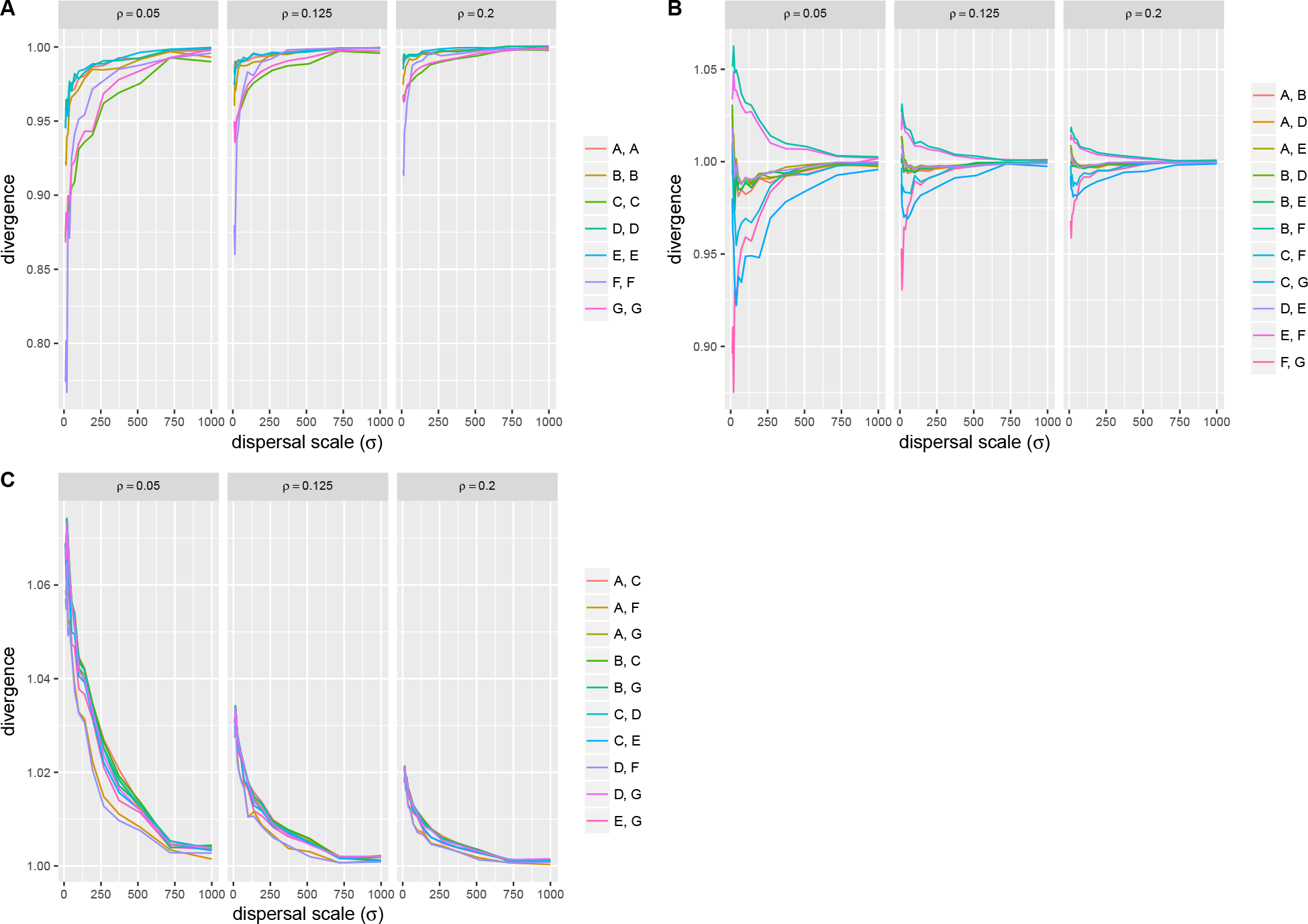
All pairwise divergences (scaled to mean divergence) for varying dispersal scale (*σ*) for comparisons (A) within regions, (B) between neighbors, and (C) between non-neighbors. Neighbors are defined based on King’s neighborhood; see Supplement C. Within each panel there are three subpanels labelled with the population density *ρ* (in individuals per hectare).

**Figure 7:**
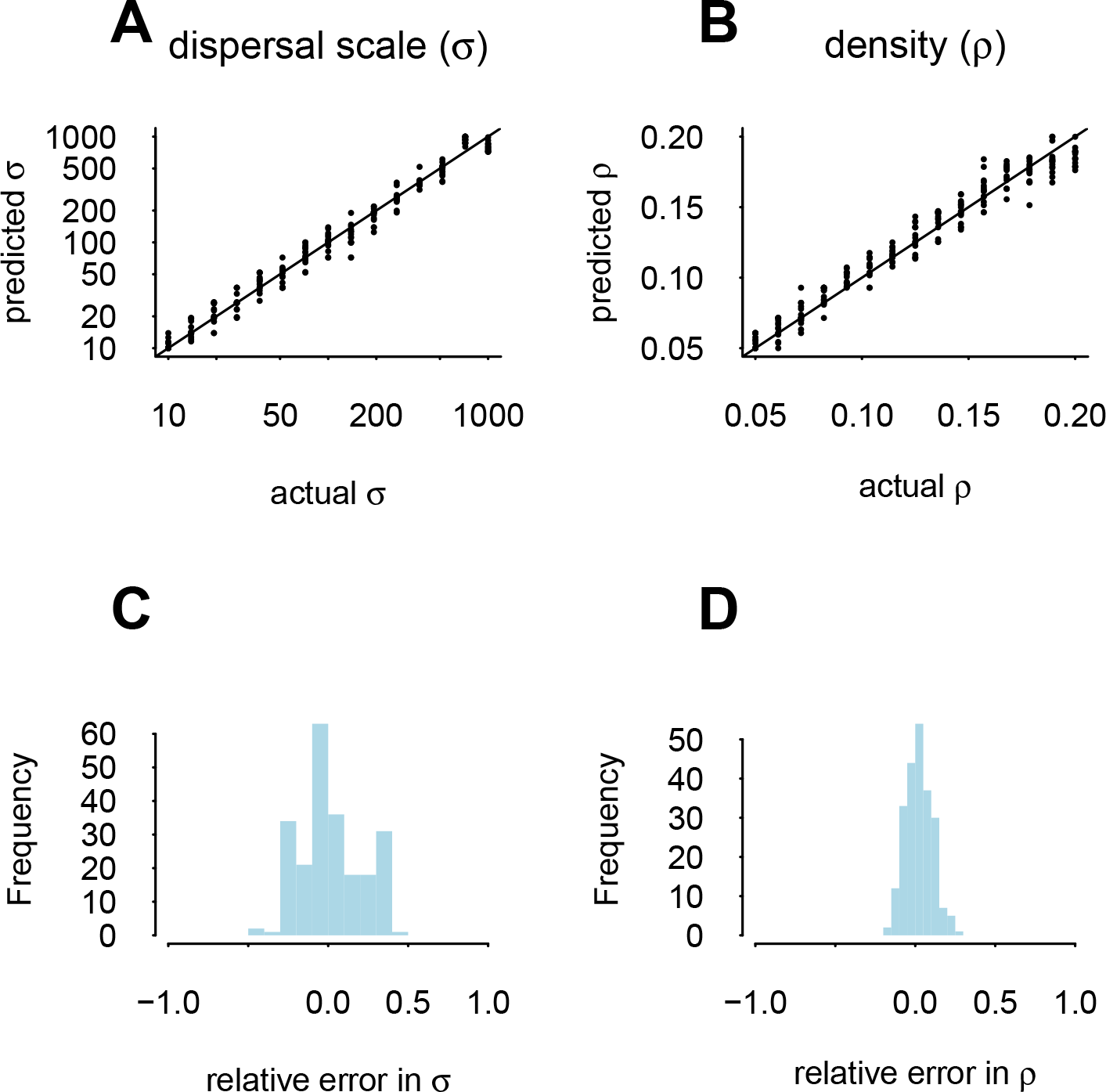
A, B) Inferred versus actual simulated parameters from 5-fold crossvalidation for dispersal (*σ*, A) and density (*ρ*, B). In each panel the solid line is 1:1. C, D) Relative error, i.e., the difference between the predicted value and the actual value divided by the actual value, for both dispersal (*σ*, C) and density (*ρ*, D).

## 4 Discussion

The methods described here enable simultaneous estimation of dispersal distance (*σ*) and carrying capacity (*ρ*) in a spatially explicit model of an age-structured population. Using data simulated from a model for which other parameters (e.g., age-specific survival) are fixed based on empirical values for *G. agassizii*, we show that our methods can estimate dispersal and carrying capacity parameters to within 10% of their true values. Thus, these methods provide a way to estimate parameters for which it is difficult to obtain data within the context of relatively well-known parameters.

In addition, we have introduced spatial analogues of common population genetic statistics and showed how and why they contain signal about geographic dynamics. Over the past decade, the four-point statistic has been widely applied in population genetics (Reich et al. 2009; Peter 2016) but its utility in continuous geography has not been appreciated. Here we show that the information derived from calculating all possible statistics for a partitioning of continuous space provides sufficient information to recover spatial population dynamics. However, as the equations in Figure 2 show, the three-point *Y* and four-point *F* are actually linear combinations of the two-point statistic *π*. Thus, to learn about geographic dynamics it would be sufficient to use only the pairwise, two-point statistic *π* in a statistical learning method that discovers correlations between outcome variables and arbitrary linear combinations of inputs. General purpose machine learning methods that are capable of such discovery include Random Forests (Breiman 2001) and techniques from deep learning. However, this assumes perfect data. Because the four-point statistic *F*_2_ is most robust to sequencing error as it is not affected by singleton mutations (and the other *F*-statistics can be written as linear combination of *F*_2_; Peter 2016), in practice it may be wise to also use all pairwise *F*_2_s as input data for inference.

We use inverse interpolation to estimate parameters from our simulations, and cross-validation to quantify the uncertainty in parameters inferred from simulated data. For empirical data, where the parameters are truly unknown, cross-validation cannot be used to quantify the parameter uncertainty. Therefore the bootstrap or jackknife, which like cross-validation are based on sampling, will be necessary to estimate uncertainty in estimates from empirical data.

These results show that integrating genomic data into structured ecological models is a feasible way to estimate parameters— in our case, carrying capacity and dispersal scale— for which it is difficult to gather sufficient data to estimate directly. There are additional caveats to keep in mind. First, although our method incorporates prior knowledge by fixing parameters based on literature values, it is not formally Bayesian. Because of this, the results here do not account for uncertainty in these estimates from the literature. Second, the model we used does not account for temporal or spatial variation in survival rates or fecundity.

Using our method requires spatially-resolved samples of individual genomes. In practice these locations only need be resolved to the scale of discrimination of the model. However, the spatial scale of discretization affects inference and additional work is needed to understand these effects. Further, the distribution of these samples on the landscape likely affects inference. Exactly how remains unclear, but because pairwise comparisons contain information about both density and space we suggest matching the distribution of samples to the population density.

As noted above, a lack of analytical results is a barrier to performing demographic inference with geographically explicit populations. We overcame this barrier with individual-based forward simulations. Another issue, however, is the possible influence of deep time: genetic variation can be influenced by both modern geography and existing variation present long before the present landscape became recognizable. To address this issue fully is beyond the goals of the current paper. However, as discussed in Kelleher et al. (2018) our simulation framework permits combining coalescent simulations in deep time with our forwards simulations. This approach (which Liu, Athanasiadis, and Weale 2008 called “sideways”) would permit analyzing multiple scenarios for deep time dynamics in tandem with a set of forward simulations as we have used here.

## Acknowledgements

We gratefully acknowledge helpful discussions, assistance with software, or feedback from Roy Averill-Murray, Linda Allison, Jerome Kelleher, Bo Peng, Dick Tracy, and Chava Weitzman. This project was funded in part by a grant from the USFWS and by the NSF (to HBS). This work benefited from access to the University of Oregon high performance computer, Talapas.

## Supplemental Information

### A Supplementary explanations of spatial statistics

We lay out the reasoning behind the statements on Figures 3 and 4 below with reference to the diagrams in Figure 2 depicting *Y*-statistics (with tips *A*, *B*, *C*) and *F*-statistics (with tips *A*, *B*, *C*, *D*). In each case we describe the labelling of the tips with a dictionary-like notation: {key : value}. We also denote the probability that the first coalescence involves individuals from groups *A* and *B*, for example, as 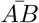.

For *Y* statistics our heuristic is 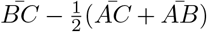. For *F*-statistics our heuristic is 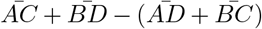. We conjecture that for both of these heuristics, if the value is positive, so is the statistic. For example, if 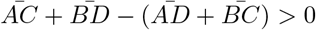 then *F* > 0.

#### A.1 Figure 3: Spatial configuration and statistics

##### A.1.1 *Y*-statistics

- i): *Y* has tips {*A*: 1, *B*: 2, *C*: 2} and because *B* = *C* then 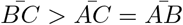 and the statistic is positive;
- ii): *Y* has tips {*A*: 1, *B*: 2, *C*: 3} and the geometry means 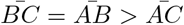 and the statistic is positive;
- iii): *Y* has tips {*A*: 2, *B* : 1, *C*: 3} and the geometry means 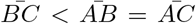 and the statistic is negative;
- iv): *Y* has tips {*A*: 1, *B* :2, *C*: 3} and the geometry means 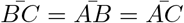 and the statistic is zero.

##### A.1.2 *F*-statistics

- i): *F* has tips {*A*: 1, *B*: 2, *C*: 1, *D*: 2} then because *A* = *D* and *B* = *C*, 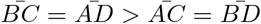 and the statistic is positive;
- ii): *F* has tips {*A*: 1, *B*: 2, *C*: 1, *D* : 3} then because *A* = *C* and the geometry, 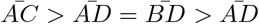 and this statistic is positive;
- iii): If *F* has tips {*A*: 1, *B*: 2, *C*: 1, *D*: 3} then because of the geometry, 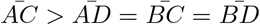 and the statistic is positive); If *F* has tips {*A*: 2, *B*: 1, *C*: 2, *D*: 3} then because 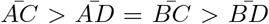 and this statistic has ambiguous sign (but if coalescences within the same group are greater than between groups, it’s positive);
- iv): if *F* has tips {*A*: 1, *B*: 3, *C*: 2, *D* : 4} then because of the geometry, 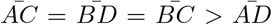, the statistic is positive; if *F* has tips {*A* : 1, *B* : 2, *C* : 3, *D* : 4} then because of the geometry, 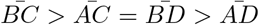 and the statistic has ambiguous sign that depends on how the chance of first coalescence involving individuals from any two groups decays with the distance between the groups;
- v): if *F* has tips {*A*: 1, *B* : 2, *C*: 4, *D* : 3} then because of the geometry, 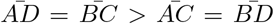, the statistic is negative; if *F* has tips {*A* : 1, *B* : 3, *C* : 2, *D* : 4} then because of the geometry, 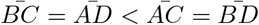 and this statistic is positive.

#### A.2 Figure 4: population size and statistics

##### A.2.1 *Y*-statistics

- i): If *Y* has tips {*A*: 1, *B*: 2, *C*: 2} and because *B* = *C* then 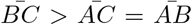 and the statistic is positive; the tips {*A*: 2, *B*: 1, *C*: 1} are equal by symmetry. When 2 is replaced with 2′, then 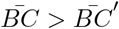 and so without other changes under the first labelling statistic must be less positive. However under the second labelling, coalescence within group 2′ has no effect and so it should be equal to the statistic on equal-sized groups.
- ii): For *Y*, coalescence within the group 1′ has no effect and so it should be equal to the statistic on equal-sized groups.
- iii): For *Y*, coalescence within the group 1′ has no effect and so it should be equal to the statistic on equal-sized groups.

##### A.2.2 *F*-statistics

- i): As in Figure 2, **line 1** but when 2 is replaced with 2′ and *F* has tips {A: 1, B: 2′,C: 1,D: 2′} then 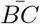 declines and the statistic is *less* positive.
- ii): As in Figure 2, **line 2** but when 1 is replaced with 1′ and *F* has tips {A: 1′, B: 2, C: 1′, D: 3} then 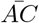 declines and the statistic is less positive.
- iii): As in Figure 2, **line 4** but when 1 is replaced with 1′ and *F* has tips {A: 1′,B: 2,C: 1′,D: 3} then 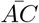 declines and the statistic is less positive.

### B Branch lengths or sequence details

Using `msprime 0.5.0` we simulated populations on a 10-subpopulations stepping stone with the total migration proportion out of each subpopulation set to 10^−4^ per generation. For each of 5 replicate populations we performed coalescent simulations of 500 samples for a genome length of 10^6^ base pairs with recombination rate and mutation rate both set to 10^−8^ per base pair. Code for the simulations is shown below. For both *F*_4_ and *Y*_3_ we chose 22 random statistics (among all possible using the 10 subpopulations as groups) and computed both using site-based and branch length-based methods. The results are shown in Figure B.1.

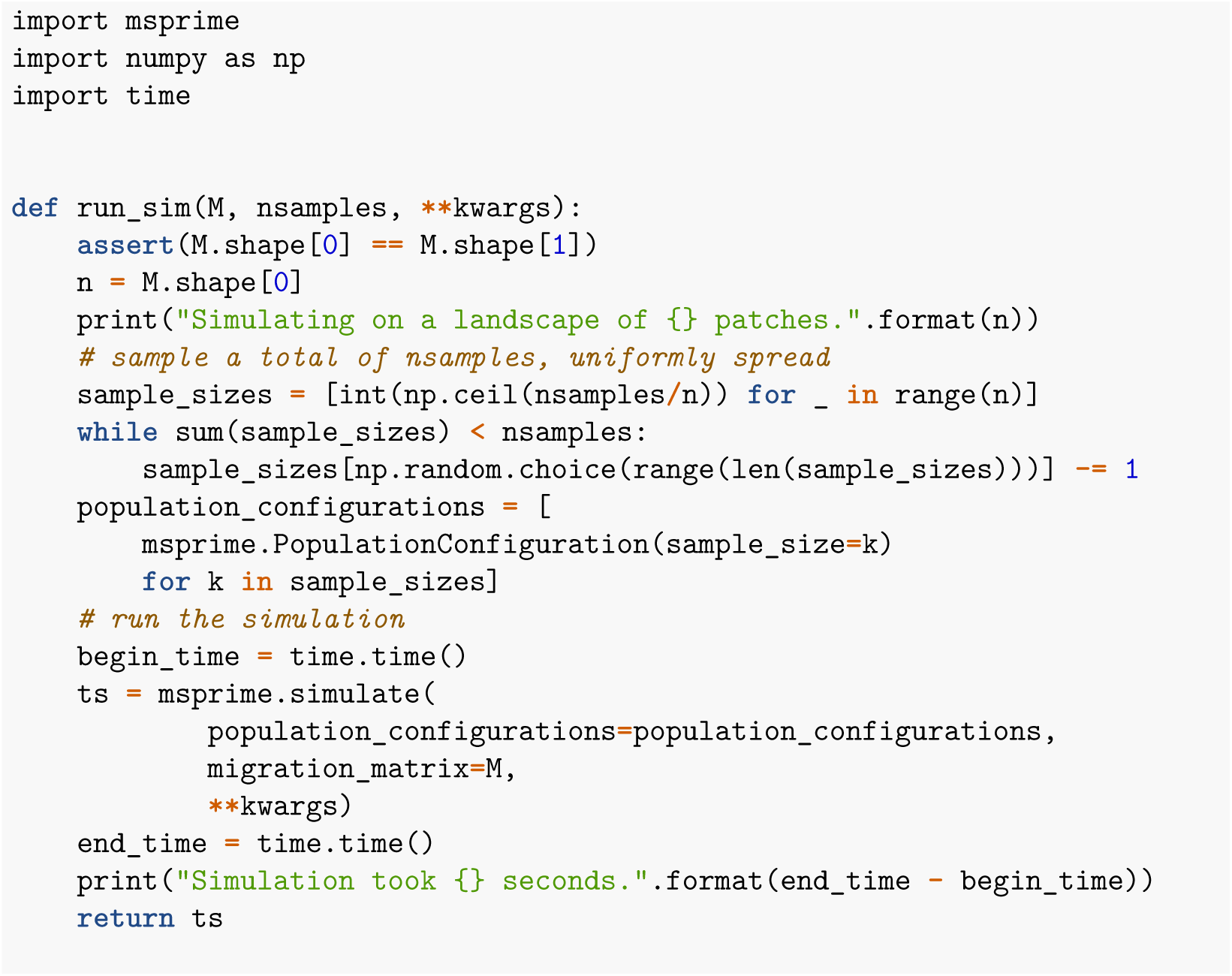
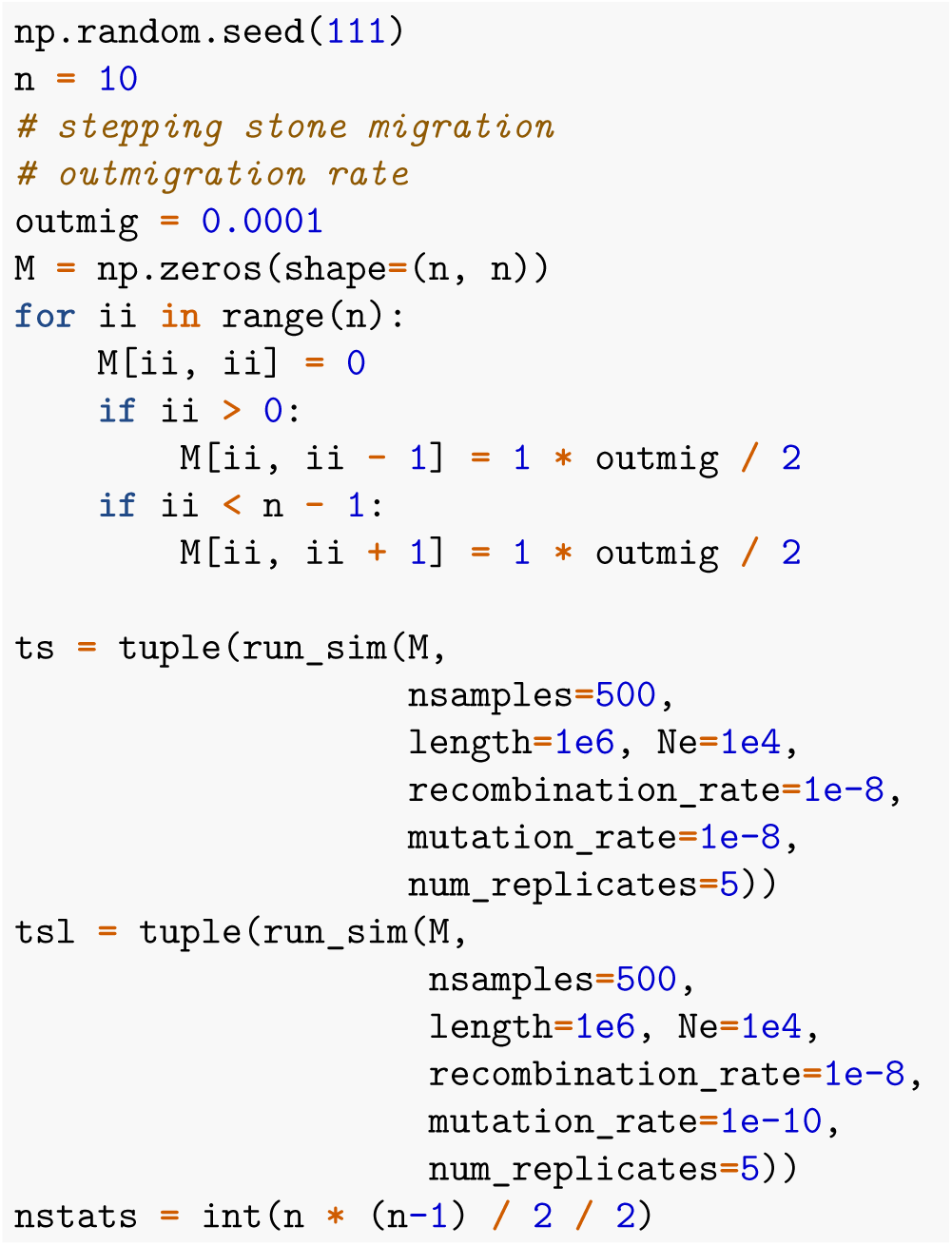

### C Biologically-motivated custom statistics

The first custom statistic aims to capture the overall timescale over which sampled alleles find a common ancestor; for this we use divergence averaged across all comparisons. The next three custom statistics aim to quantify the timescale over which individual alleles sampled from opposite sides of the population range find a common ancestor; for this we apply the three-point statistics to groups spanning the range, with the focal group in one corner and the two comparison groups on the opposite end of the landscape. For example, the *Y* statistic with a focal group in the northwest corner and the comparison groups on the east side is *Y*_3_(*A*; *C*, *G*). Specifically, we computed *Y*_3_ with focal groups in the west, 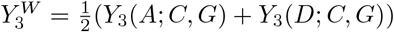; and east 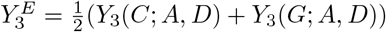. We also computed 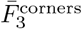, the average of the four *F*_3_ statistics having a corner population as focal and the two most distant corner populations as the other two arguments.

The final three custom statistics aim to quantify differences in timescales over which alleles find common ancestors depending on whether they are sampled from i) within the same region, ii) neighboring regions, or iii) non-neighboring regions. To do this, we categorized every divergence statistic *π*(*A*, *B*) according to these three categories, i.e., whether i) *A* = *B*, ii) *A* and *B* were neighbors, or iii) otherwise, and averaged mean divergences within each of these three categories, denoting these quantities *π*_*w*_, *π*_*n*_, and *π*_*nn*_, respectively. Then, we took differences of these average divergences to create three statistics: between neighbors and within-region, *π*_*n*_ — *π*_*w*_; between non-neighbors and within-region, *π*_*nn*_ — *π*_*w*_; and between neighbors and non-neighbors, n_*n*_ — *π*_*nn*_. The neighbors and non-neighbors for each group in Figure 1C (using a King’s neighborhood) are listed in Table C.1.

We compare performance of interpolating parameter values from two different sets of statistics. First, all the statistics meaning every one of the six types of statistics computed across all 7 groups (*n* = 406). Second, **custom** statistics only, meaning the biologically motivated predictors (*n* = 7): mean divergence 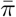, neighbor vs within-region divergence *π*_*n*_ − *π*_*w*_, non-neighbor vs within-region divergence *π*_*nn*_ − *π*_*s*_, non-neighbor vs neighbor divergence *π*_*nn*_ − *π*_*n*_, 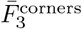, 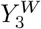, and 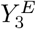.

**Figure B.1:**
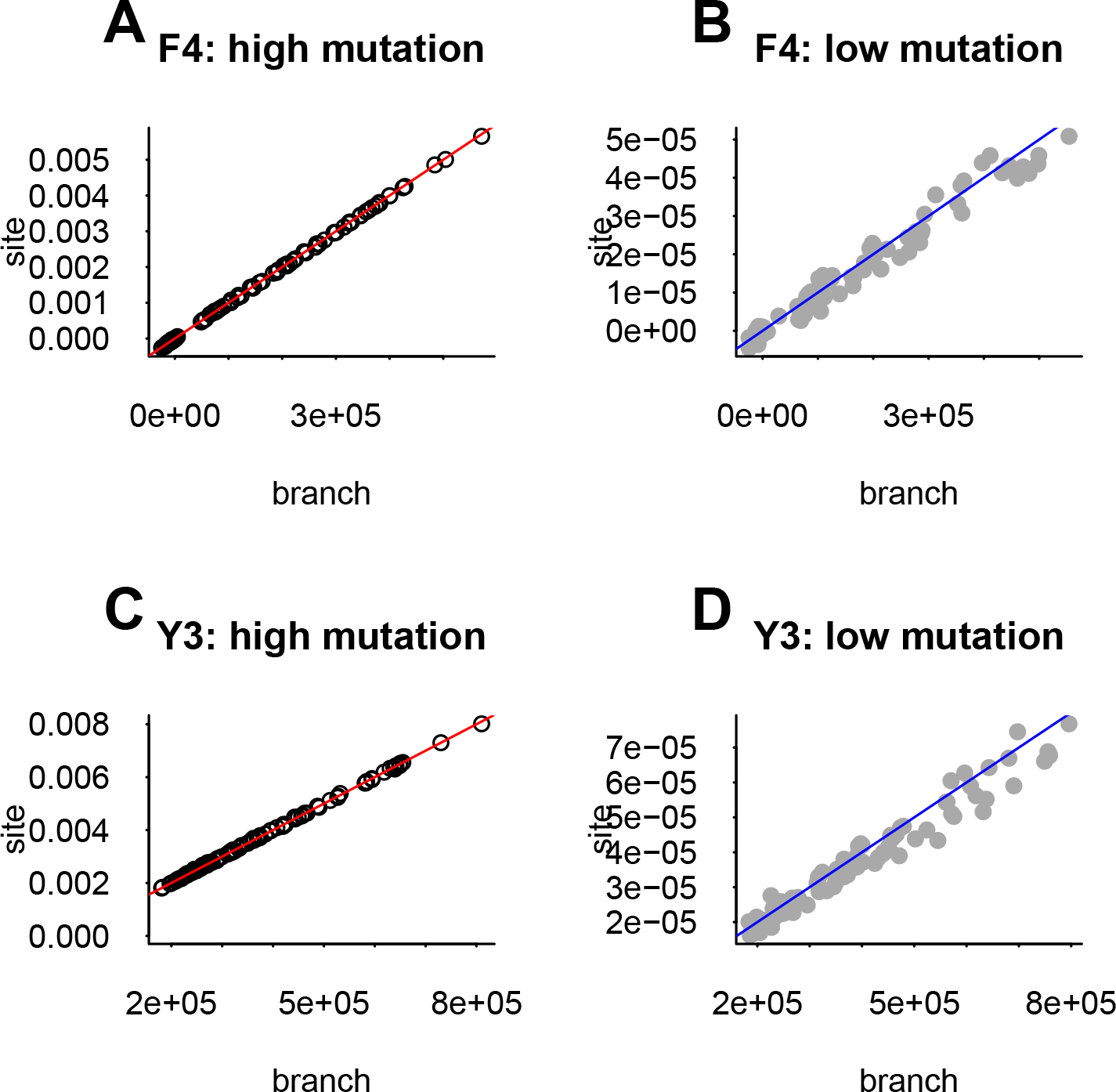
Comparing *F*_4_ and *Y*_3_ calculated using sequence data (y-axes) and using branch lengths of marginal genealogies (x-axes) for high (circles and red lines) and low (gray dots and blue lines) mutations rates. The slopes of the lines in each plot are the mutation rates.

**Table C.1:** Regions, their neighbors, and their non-neighbors.

| region | neighboring regions | non-neighboring regions |
| --- | --- | --- |
| A | B, D, E | C, F, G |
| B | A, D, E, F | C, G |
| C | G | A, B, D, E, F |
| D | A, B, E | C, F, G |
| E | A, B, D, F | C, G |
| F | B, E, F, G | A, C, D |
| G | C, F | A, B, D, E |

#### Intuition from custom statistics

For example, the difference between 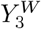 and 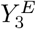 likely relates to the relative strength of migration and population sizes on the east and west parts of the landscape. Between subpopulation migration is higher and population sizes are lower on the eastern part of the landscape. This means that when the focal group is in the west, the other two groups are likely to have coalesced earlier and thus *Y*_3_ has a larger value. This is similar to Figure 4i where the three-point statistic decreases in magnitude when the non-focal population is larger: *Y* (1; 2, 2) > *Y* (1; 2′, 2′) because group 2′ has a larger population size than 2.

### D Simulation parameters

Simulations were run for 15000 years, starting from the age distribution shown in Figure D.3, which is roughly at equilibrium. Table D.2 gives age-specific survival and fecundity. Genetics were specified as a single chromosome of length of 10^8^ base pairs with 10 recombining loci and a recombination rate of 10^−6^ per generation.

**Table D.2:**
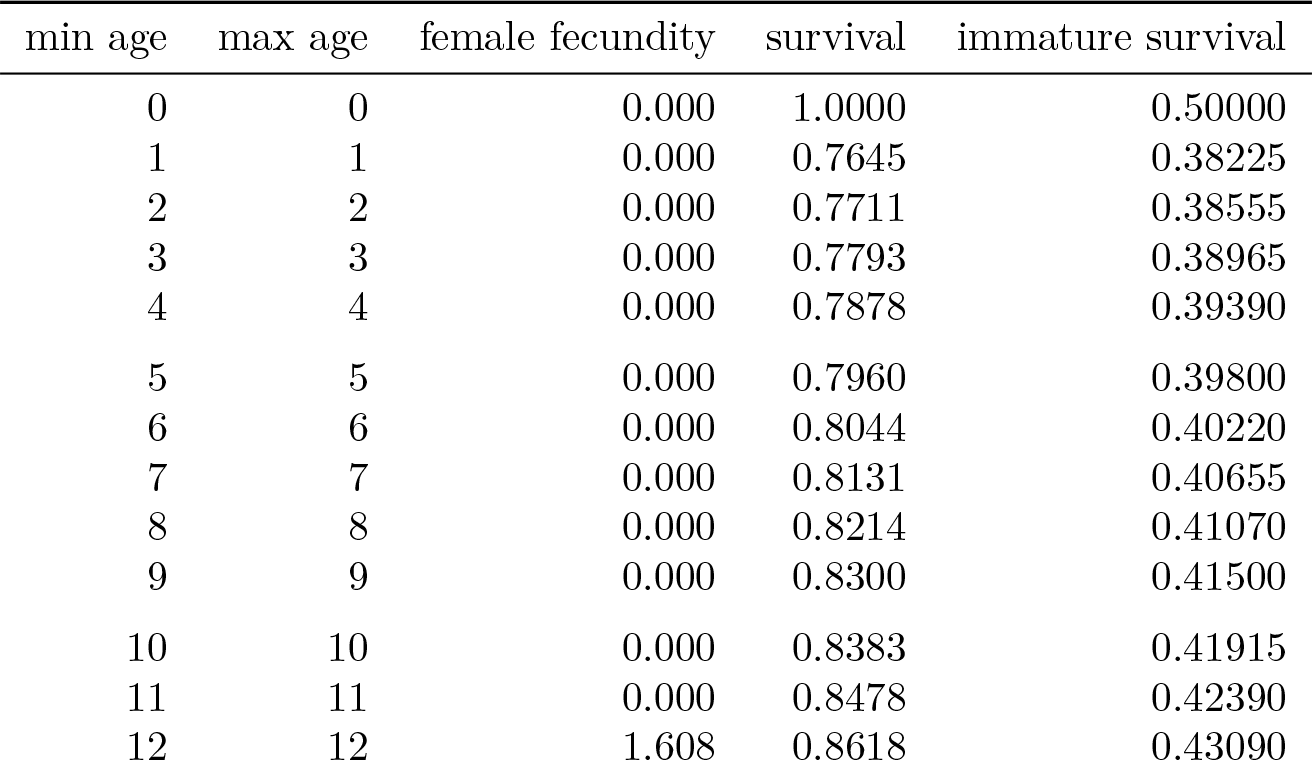
Life table of age-based female fecundity and survival after Reed *et al*. (2009) but with fecundity (*r*_0_) doubled (life tables model only female offspring). Males are non-reproductive below age 15 and of equal fitness above this cutoff. Immature individuals survive at 0.5 the rate of immature individuals.

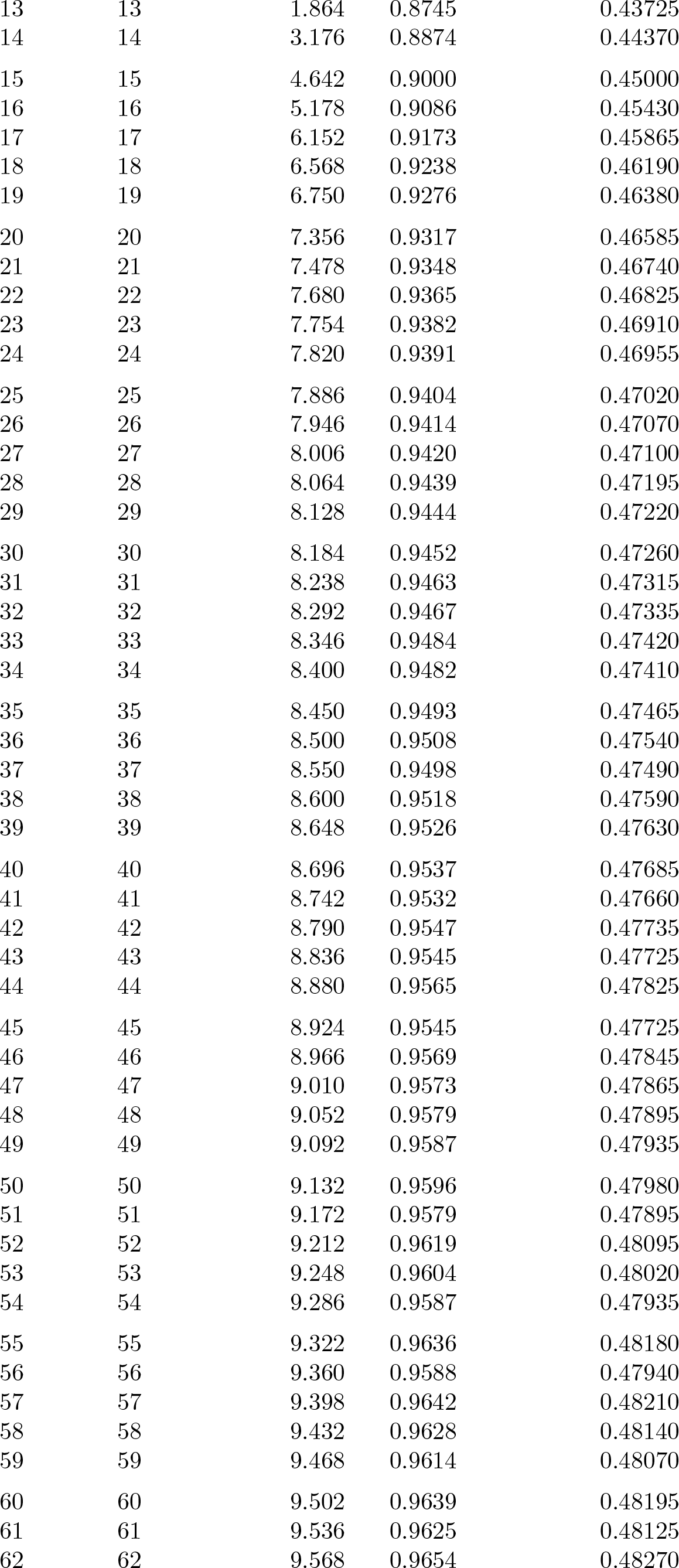

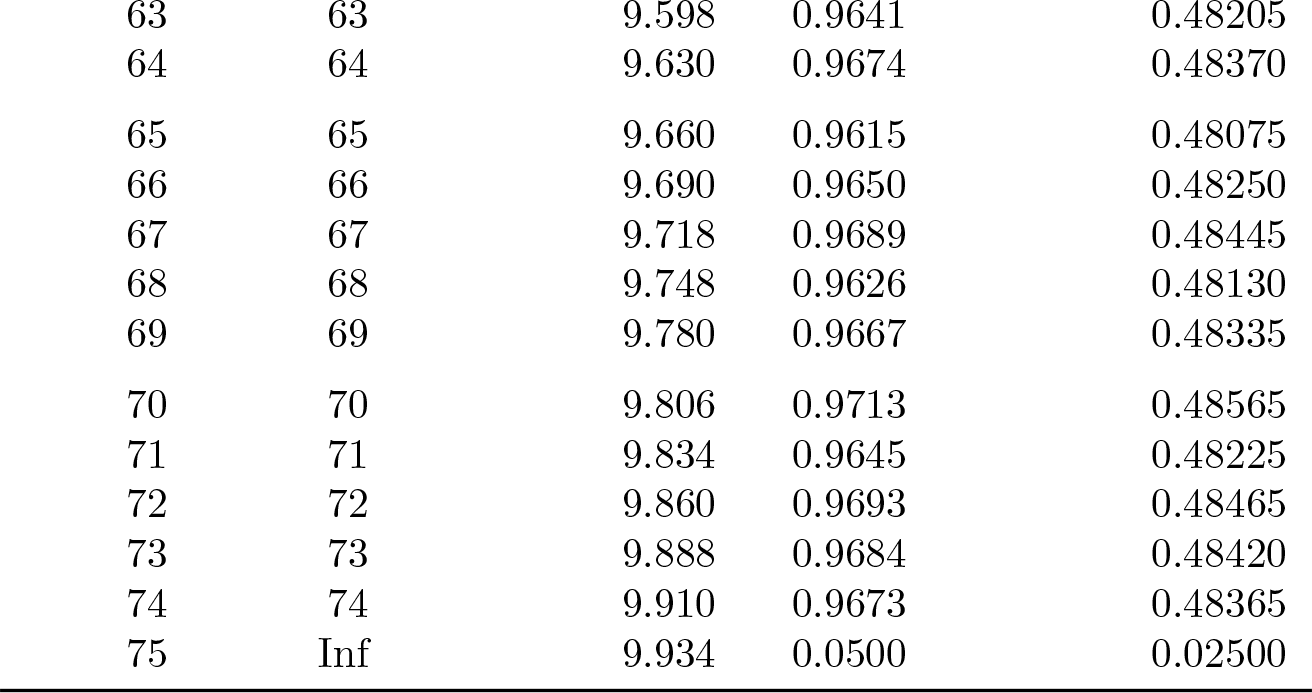

The mean realized lifetime fitness (number of offspring) versus age for both males and females is shown in Figure D.4. This quantity is computed from simulated data by sweeping forward in time and recording the number of offspring produced after an individual reaches a given age (recorded on the x-axis); the y-axis shows the mean of this quantity versus age.

### E Extended Results

#### E.1 All statistics

With all possible statistics shown, the biological meaning in the patterns is difficult to discern (Figure E.5).

#### E.2 Crossvalidation results

Figure E.6 shows the median relative error (black dots) across all k folds of cross-validation for our method, either with “custom” statistics or all possible statistics. In all cases, the method inferred parameters within a few percent of their true (simulated) values. The number of folds in cross-validation did not much affect performance.

**Figure C.2:**
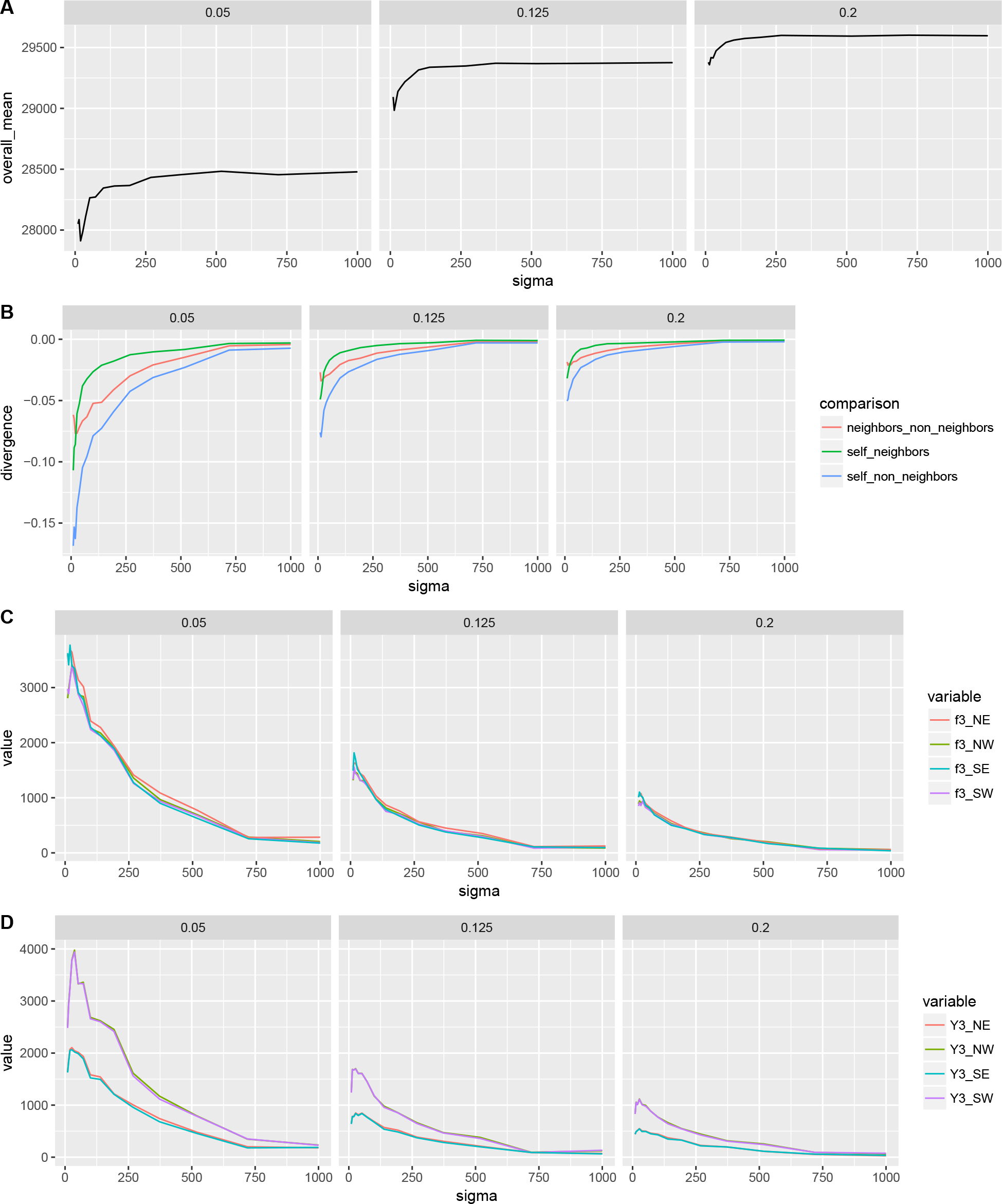
Custom statistics for varying dispersal scale (*σ*). Within each panel there are three subpanels labelled with the population density ρ in individuals per hectare (0.05, 0.125, 0.2). A) Divergence averaged across all populations. B) divergence_*n*−*nn*_ scaled by mean divergence for neighbors defined in Table C.1. Three-point statistics with focal population in NW, SW, SE or NE corner (see legend): C) *F*_3_, in the main text 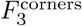 is the average of the lines shown, and D) *Y*_3_, in the main text 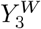 is an average of the values for W focal populations and 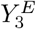 is an average of the values for E focal populations.

**Figure D.3:**
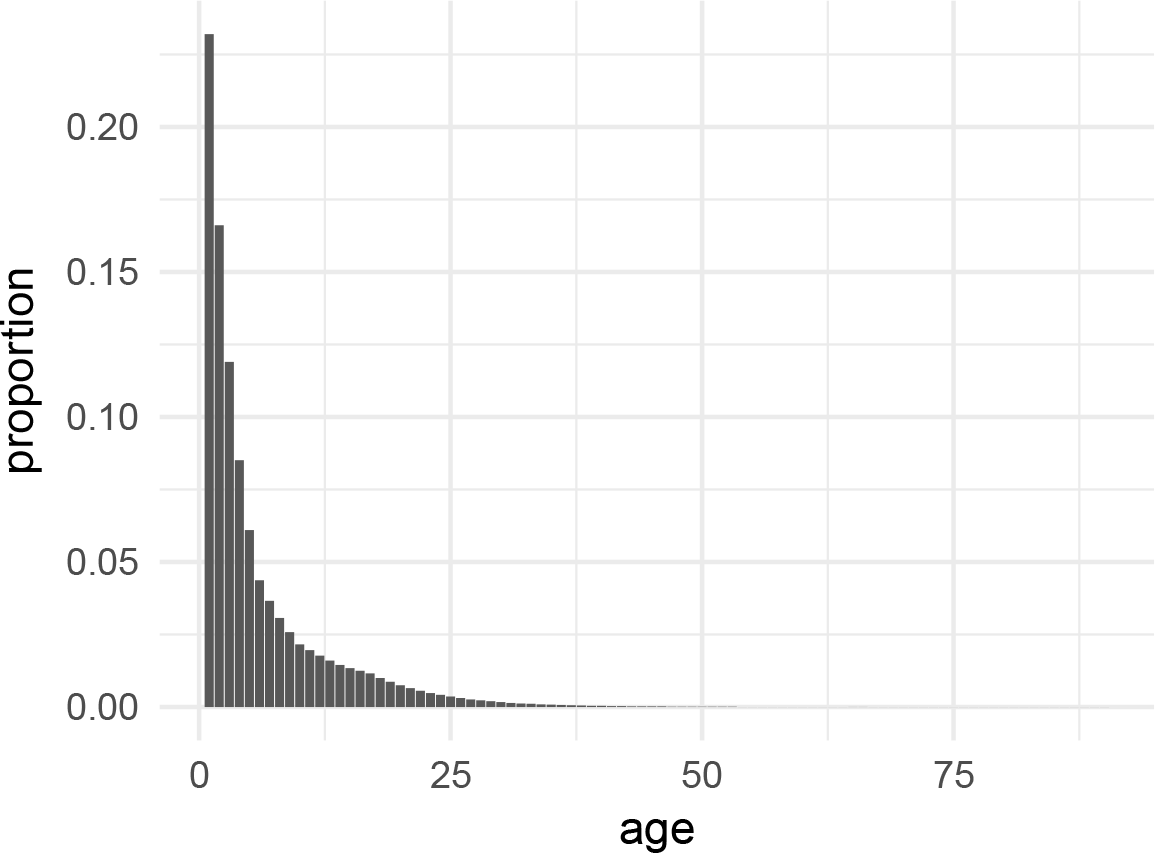
Initial age distribution

**Figure D.4:**
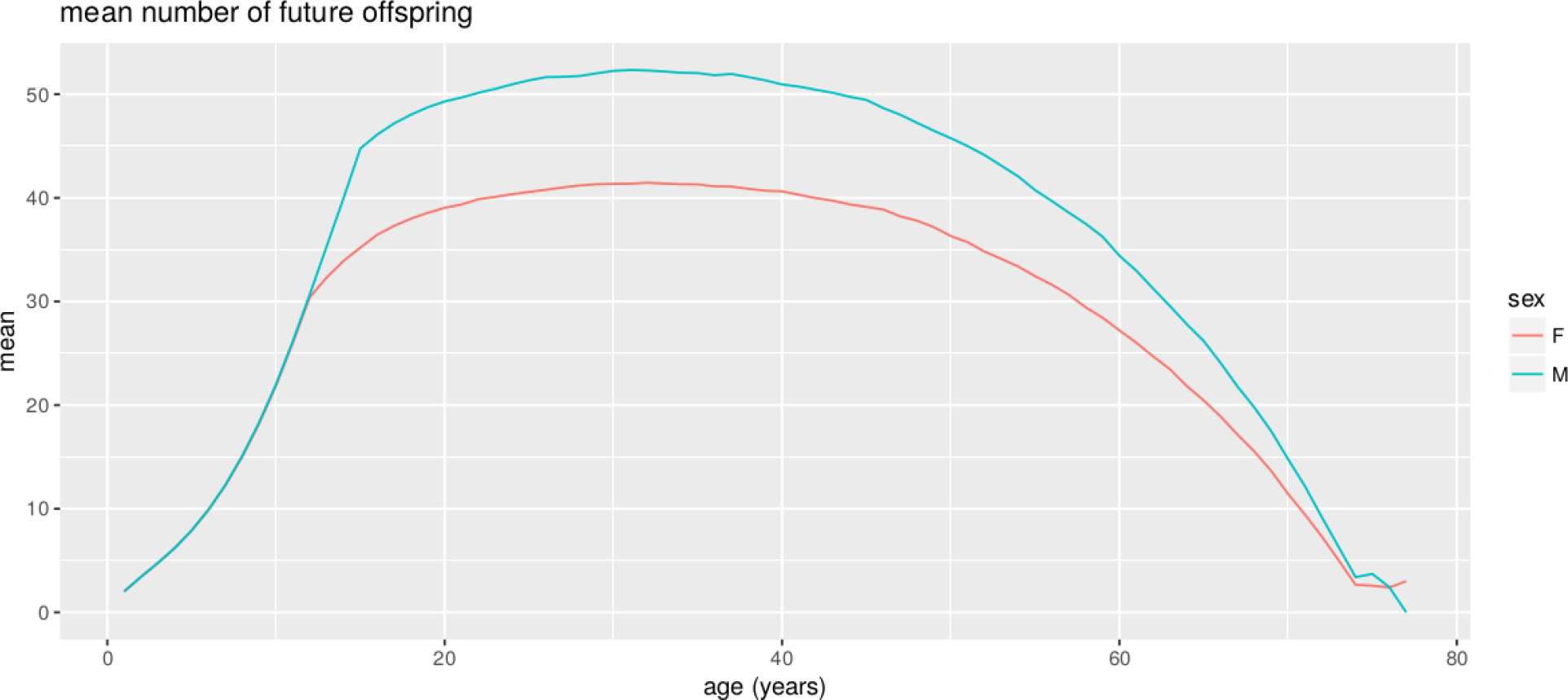
Mean number of future offspring versus age (analog of reproductive value) observed in the model.

**Figure E.5:**
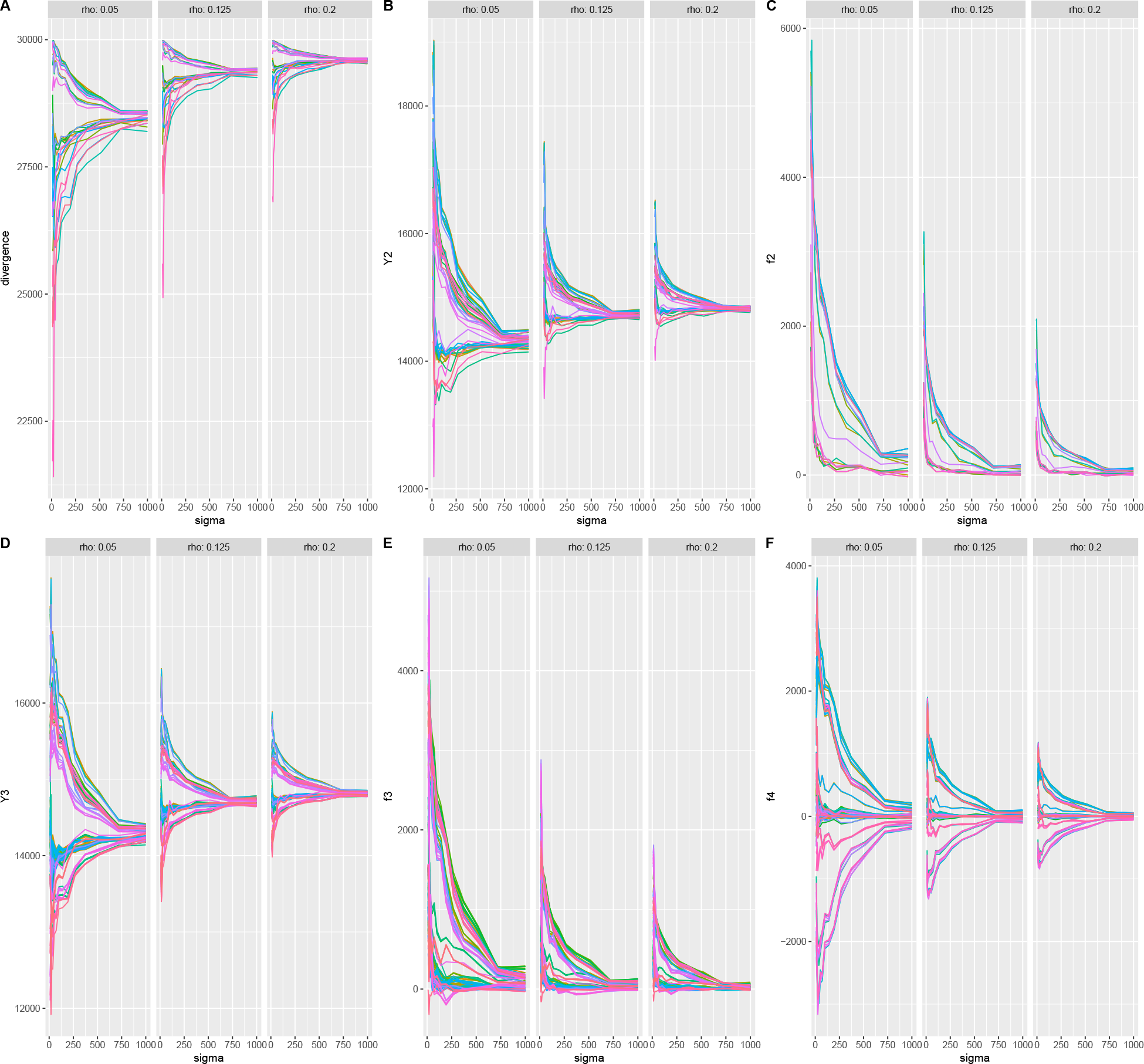
All values of (A) divergence, (B) *Y*_2_, (C) *F*_2_, (D) *y*_3_ (recall *Y*_3_(*A*; *B*, *C*) = *y*_3_(*a*; *b*, *c*) — 1/2(*y*_3_(*b*; *a*, *c*)+ *y*_3_(*c*; *a*, *b*))), (E) *F*_3_, and (F) *F*_4_ across all combinations of the 7 groups for varying dispersal scale *σ*. Within each panel there are three subpanels labelled with the population density *ρ* in individuals per hectare (0.05, 0.125, 0.2). Note differeing *y*-axis scales.

**Figure E.6:**
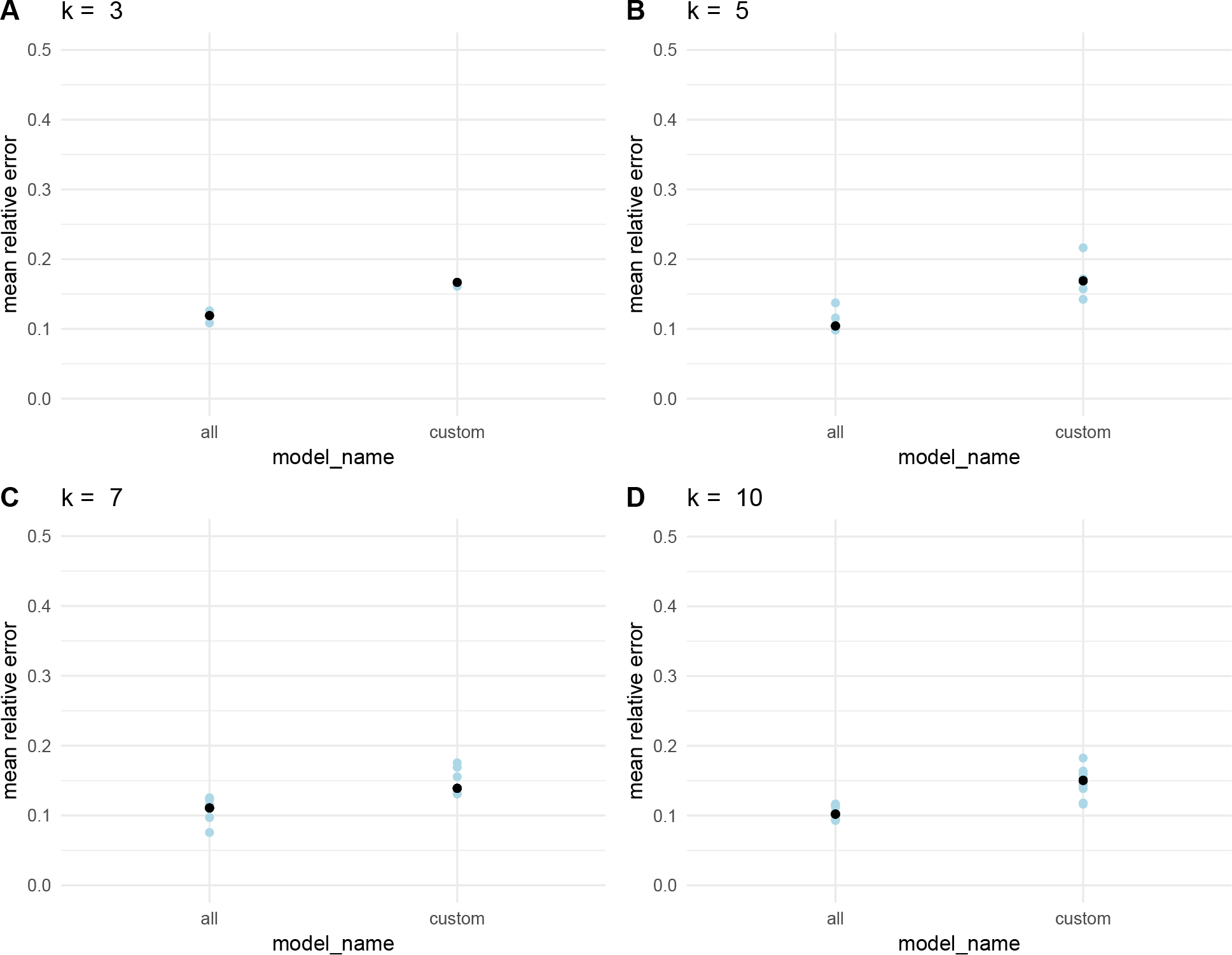
Model performance under *k*-fold crossvalidation (for values of k noted in the titles) for both inverse interpolation with all predictors or just a few biologically motivated ones: mean relative error (blue dots) of the *k* crossvalidation replicates and the median across all replicates (black dot).

